# Eco-evolutionary dynamics of active virus-host interactions in a freshwater lake: revealed through metaHi-C

**DOI:** 10.64898/2026.02.18.706586

**Authors:** Francisco Nadal-Molero, Ana-Belen Martin-Cuadrado, Maliheh Mehrshad

## Abstract

Detecting active phage–bacteria interactions in natural microbial communities remains a major limitation for understanding their ecological dynamics and associated co-evolutionary processes. Here, we applied metaHi-C, a chromosome conformation capture method, to resolve active virus–host associations in a freshwater microbial community. From >900 microbial and >33,000 viral Hi-C–assembled genomes, we identified 100 high-confidence phage–host linkages spanning major freshwater bacterial lineages, including *Limnohabitans*, *Acidimicrobium*, *Synechococcus*, *Candidatus* Nanopelagicus, *Candidatus* Planktophila, *Candidatus* Methylopumilus and *Polynucleobacter*. The inferred networks revealed diverse infection patterns, including broad-host-range phages, cellular-level co-infection, kill-the-winner dynamics and one-to-one interactions. These ecological patterns were associated with signatures of diversifying selection in host-interaction genes, consistent with host-range expansion, alongside conserved genomic regions in broad-host-range and co-infecting phages, indicating functional constraints on essential infection modules. Together, these results demonstrate that metaHi-C enables direct linking of community-level infection dynamics to underlying evolutionary processes, revealing how these forces shape bacterial population dynamics of freshwater bacteria.

## INTRODUCTION

Dependence of phages on their bacterial hosts for replication has defined the evolutionary trajectories and ecological dynamics of both parties for millions of years. However, disentangling their ecological interactions at the ecosystem level remains a major bottleneck, hindering our understanding of phage-bacteria coevolution. Phages are ubiquitous and ecologically essential due to their important role in shaping bacterial community dynamics, gene flow, and ecosystem-level nutrient cycling^1–3^. Large-scale environmental metagenomics studies have significantly improved our understanding of phage genomic diversity and evolution; however, they are inefficient at resolving phage-bacteria interactions^4–6^. Cultivation, the gold standard for studying phage-bacteria interactions for many years, suffers from the “great plate count anomaly”^7^, and since only a small fraction of environmental microbes can be maintained in the laboratory^8^, the majority of our knowledge about phages stems from those infecting a narrow set of model bacterial hosts. As a result, the full spectrum of virus-host interactions in natural samples remains vastly underexplored. For instance, ∼90% of viral sequences recovered from aquatic metagenomes are still unclassified and unlinked to hosts^3^, reflecting the severe lack of direct data on virus infection dynamics and host interactions at the ecosystem level.

Freshwater environments exemplify these blind spots, where bacterial community is dominated by fastidious lineages (e.g., Actinobacteria^9,10^, streamlined methylotrophs^11^, or members of the LD12 clade of Alphaproteobacteria^12^ that have eluded cultivation for decades, resulting in a dearth of phage isolates^13^. Compounding this challenge is the difficulty of linking uncultivated phages to their natural hosts in metagenomic and metaviromic studies^14^. Inference-based *in-silico* methods, including CRISPR spacer matching^15,16^, oligofrequency similarity^14,17,18^, tRNA homology^19^, sequence composition models^20^, machine learning approaches^21^, integrative frameworks like iPHoP^22^, and novel strategies leveraging shared mobile genetic elements such as MITEs^23,24^, have been used for host prediction. Yet these methods often yield overly broad taxonomic assignments, and even when successful, they apply to only a small fraction of the phage diversity and rarely confirm active infections.

Environmental evidence increasingly challenges the traditional view that viruses typically exhibit strict host specificity, being restricted to one or a few closely related host species^25^. Culture-independent studies suggest that broad-host-range phages may be more prevalent than previously assumed^26^, and that viruses can engage in cryptic infections of non-primary hosts, establish latent infections, contribute genetic material without triggering lysis^27^, or undergo host-switching events not captured by standard assays^6,28,29^. This suggests that the view of viruses as one-shot killers that infect and lyse a single, specific host, is overly simplistic and should be expanded to a more nuanced and flexible model of infection, particularly in dense microbial communities with poorly defined host boundaries^30^.

To inspect phage infection dynamics, direct detection of individual phage-host associations is required. Unlike inference-based *in-silico* host assignment methods, high-throughput chromosome conformation capture (metaHi-C) has recently emerged as a powerful technique for detecting ongoing/active infections by linking DNA interactions between phages and their host genomes in mixed communities^31,32^. Here, we apply metaHi-C to a freshwater microbial community and capture a snapshot of ongoing infections, reconstructing the active infection network that underpins the functioning of this aquatic ecosystem. MetaHi-C results provide direct evidence for the prevalence of broad-host-range phages, instances of intense infection pressure from multiple phages targeting the same host population, and co-infections involving two phages within a single bacterial cell. These findings reveal that such events are not rare anomalies but ecologically relevant phenomena with significant impact on freshwater microbiome. Our results challenge the assumption of one-virus-one-host specificity and dynamics in natural ecosystems and reveal the complexity of viral ecology with vast implications for horizontal gene transfer, viral competition, and host eco-evolutionary dynamics.

## RESULTS AND DISCUSSION

### Reconstructing freshwater community using MetaHi-C

A freshwater sample was collected in July 2024 from the Amadorio reservoir (Alicante, Spain) and used to generate two proximity ligation libraries, which were then merged into a single metaHi-C dataset, AmaJul24-HiC (34 Gb, Illumina PE250). A reference metagenome was also generated from the same sample, AmaJul24 (80 Gb, Illumina PE150).

Binning the assembled contigs of AmaJul24 (380,079 contigs >1 kb) using the metaHi-C derived linkages resulted in 1043 Hi-C assembled genomes, with only 25 exceeding the 5% contamination threshold in accordance with MIMAG standards for high-quality reconstructed genomes^33^ (Supplementary Figure S1a). In parallel, 38,602 contigs were identified as viral and binned in viral Hi-C-assembled genomes (vHAGs) (Supplementary Figure S1b), with only 96 classified as complete based on CheckV^34^. The small genome sizes of reconstructed HAGs (median of 122.1 kb and ranging from 30 kb to 7.2 Mb) hint at over-fragmentation as a potential cause of the HAG incompleteness (only 4% of HAGs had completeness ≥50%, Supplementary Figure S1a).

To test whether linkage-based binning leads to genome over-fragmentation, the same method was applied to a Hi-C dataset generated from a reference infection model consisting of an axenic *Escherichia coli* K12 culture and its bacteriophage Lambda. Using the same workflow, the *E. coli* genome was fragmented into five different genomic bins (Supplementary Figure S2). This over-fragmentation can be attributed to the 3D conformation of prokaryotic DNA into so-called “Chromosomal Interaction Domains” (CIDs), which, as in eukaryotes^35^, leads to an uneven distribution of linkages along the genome and the consequent misinterpretation of densely linked regions as separate bins. In this case, the Hi-C contigs align with the known CIDs of *E. coli*^36,37^, supporting the validity of our hypothesis. To address potential over-fragmentation in the linkage-based binning of AmaJul24, the reconstructed HAGs/vHAGs were manually curated using a custom metric - the *Nadal Index* (NI) (see Methods). This index quantifies Hi-C linkage density within and between pairs of HAGs, identifying candidates for merging as fragments of the same genome. Visual inspection of the Hi-C contact maps for each proposed merge confirmed their relatedness as parts of the same original genome. Applying the NI, followed by manual curation, identified 176 HAGs (17% of the total) as candidates for merging, reducing the number of HAGs from 1,043 to 903 (Supplementary Table S1). This merging increased completeness for 36 joined HAGs (median rising from 7.1% to 80.4%) with minimal or no increase in their contamination scores. In parallel, 13 vHAGs were also identified for merging (Supplementary Table S1).

Reconstructed HAGs were dominated by representatives of ubiquitous freshwater lineages, including *Acidimicrobium* (*n*=93, 8.92%), *Limnohabitans* (*n*=86, 8.25%), *Ca.* Planktophila (*n*=63, 6.04%) and *Ca.* Nanopelagicus (*n*=57, 5.47%) (Supplementary Figure S3, Supplementary Table S2). The majority of vHAGs were affiliated to Caudoviricetes (*n*=23,937, 70.78%) and Megaviricetes (*n*=2,872, 8.49%), with 6,706 vHAGS (19.83%) remaining unclassified (Supplementary Table S3). Clustering vHAGs at 95% average nucleotide identity (ANI) over 85% of the genome length resulted in 33,394 viral operational taxonomic units (vOTUs).

### Hi-C linkages revealed active infection of major freshwater bacterial lineages

Hi-C linkage data were utilized to identify active virus-bacteria interactions within AmaJul24 dataset. Analyses of curated linkages revealed 100 high-confidence virus-host associations involving 85 vHAGs and 73 HAGs (Figure 1a&b). Within this network, HAGs affiliated with the genus *Limnohabitans* were the most prevalent (n=7, 9.59%), followed by other freshwater lineages including *Acidimicrobium, Synechococcus*, *Ca.* Nanopelagicus, *Ca.* Planktophila (each n=5, 6.85%), and *Ca.* Methylopumilus (n=4, 5.48%) (Figure 1c & Supplementary Table S4). A comparison of coverage-derived abundances (based on read mapping) between linked *vs* unlinked vHAGs in AmaJul24 metagenome revealed that linked vHAGs were significantly more abundant (Wilcoxon test = 3,203,820, *p* < 2.2·10^-16^; Supplementary Figure S4a). A similar pattern was observed among HAGs, where linked HAGs also displayed moderately higher coverage compared to unlinked counterparts (Wilcoxon test= 40,654, *p* = 1.4·10^-7^; Supplementary Figure S4b). These results highlight that abundant bacterial lineages are more likely to be actively infected by viruses, and that Hi-C has higher resolution for detecting interactions in dominant taxa, an effect enhanced by the stringent criteria applied for linkage validation.

**Figure 1.**
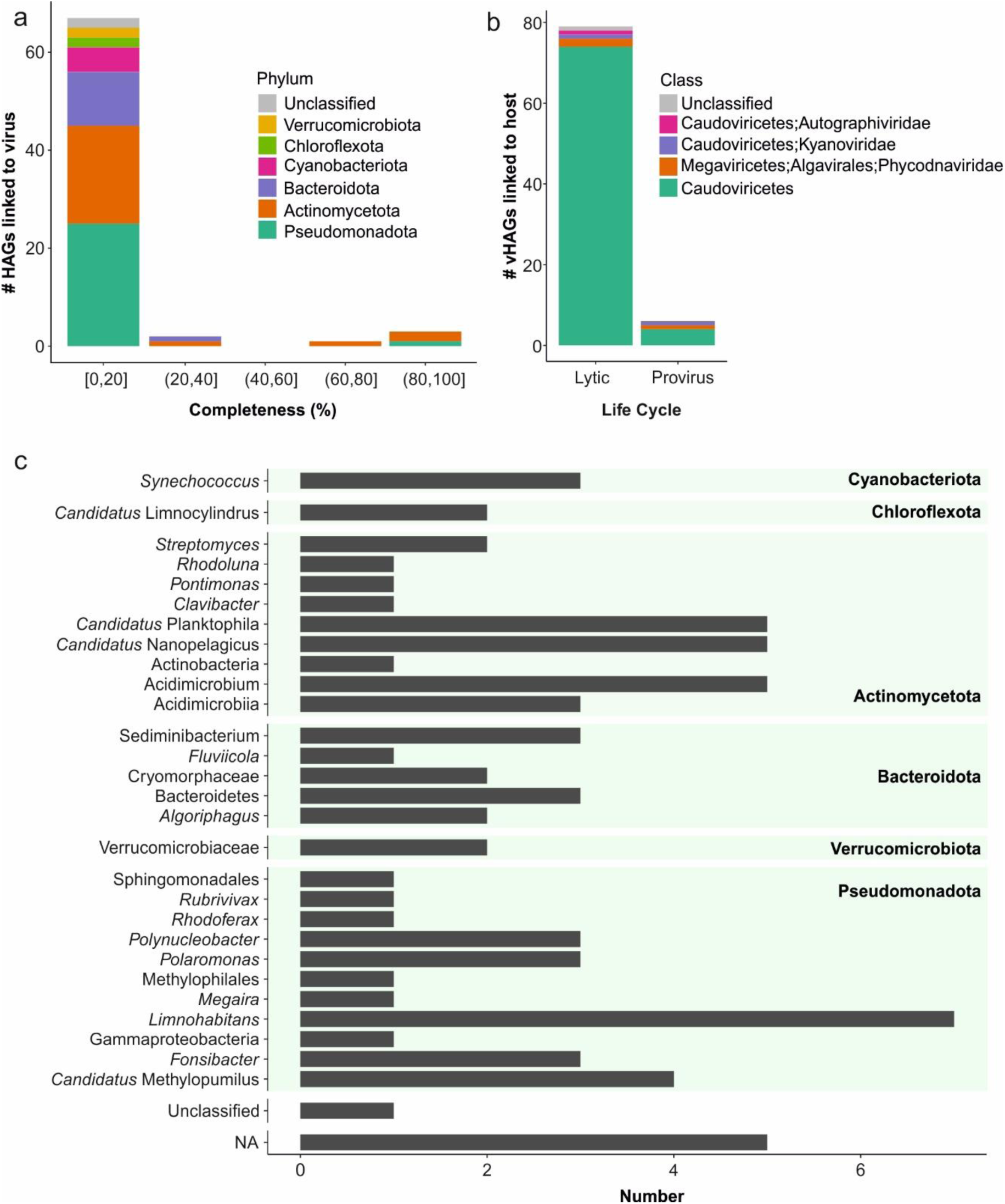
Taxonomy of linked HAGs and vHAGs in the AmaJul24 dataset based on Hi-C linkages. (a) Phylum-level taxonomy of linked HAGs across different completeness spectrum (b) Taxonomy of linked vHAGs grouped by their predicted life cycle. (c) Higher- resolution taxonomy of HAGs linked to at least one vHAG.

Notably, the most abundant HAG in our dataset (24,390 rpkgs), affiliated with the filamentous Cyanobacteria *Aphanizomenon,* lacked any detectable link to a vHAG (Supplementary Table S2). This HAGs high coverage may result from a bloom, with cells covered by a mucilaginous sheath^38^ that hampers phage access, or from the presence of akinetes, dormant cells with minimal metabolic activity and high polyploidy^39^, which are likely to be highly resistant to phage infection. The second most abundant HAG, classified as Bacteroidota (24,270 rpkgs), was linked to vHAG0025.

Based on CheckV predictions, both lytic (n=79) and lysogenic (n=6) vHAGs were found to be linked to hosts, and virus-per-host (VPH) estimates showed slightly higher relative abundances for lytic viruses compared to lysogenic ones, as expected (Supplementary Figure S5). The majority of linked vHAGs belonged to class Caudoviricetes (n=81, 95.3%), while three were classified as Megaviricetes (3.6%), and one vHAG remained unclassified (Figure 1b). The most abundant vHAGs were both classified as Caudoviricetes (75,060 and 61,671 rpkgs, respectively), yet neither was linked to a host. This could be due to rapid degradation of host DNA during the later stages of phage infection or, alternatively, that these vHAGs belong to phages captured in the final stage of infection, with capsids already formed and their DNA highly packaged^40,41^. The next four most abundant vHAGs (rpkgs values ranging from 38,799 to 31,444) were linked to a Verrucomicrobiaceae and the other three to *Synechococcus-*affiliated HAGs.

Most vHAG-HAG associations followed a 1:1 pattern (n=41); however, 11 vHAGs were associated with multiple HAGs (up to four), representing broad-host-range phages (Figure 2). Conversely, in two particularly striking cases, *Synechococcus* HAGs, HAG5 and HAG8, were linked to 14 and 6 distinct vHAGs, respectively, suggesting strong phage-mediated control shaping the population structure of these cyanobacterial lineages. Eight vHAGs were linked to *Limnohabitans* representatives, six to members of the genus *Ca.* Fonsibacter, four to *Ca.* Methylopumilus, and another four to *Ca.* Planktophila representatives (Figure1c & 2). None of these vHAGs showed sequence similarity to the few phages previously described for these taxa^42^; however, they show sequence similarity to sequences in the viral IMG/VR 4.1 database for which no hosts have been designated (Supplementary Information). HAGs affiliated with genus *Ca.* Nanopelagicus and *Polynucleobacter* were linked to four and three vHAGs, respectively, representing the first reported phage genomes infecting these ubiquitous freshwater lineages. Our results show that the most abundant and ubiquitous freshwater bacterial lineages are actively infected by viruses, thereby broadening the known diversity of phages infecting these lineages and resolving host associations for similar publicly available phage genomes that previously lacked host designation. To validate virus–host linkages inferred from Hi-C, we applied complementary in silico predictions. iPHoP recovered 28 host assignments (33%), with limited agreement (18%) to Hi-C results. The Actinobacteria-specific *whiB* genes (Ghai, Mehrshad et al. 2017) supported four Hi-C derived Actinomycetota-virus links, while CRISPR analysis revealed 27 matches (one corroborated by Hi-C). MITE-based screening (Nadal-Molero, Rosselli et al. 2024) yielded three additional candidate associations, but none were confirmed by Hi-C (more details in Supplementary Information & Supplementary Tables S5–S7).

**Figure 2.**
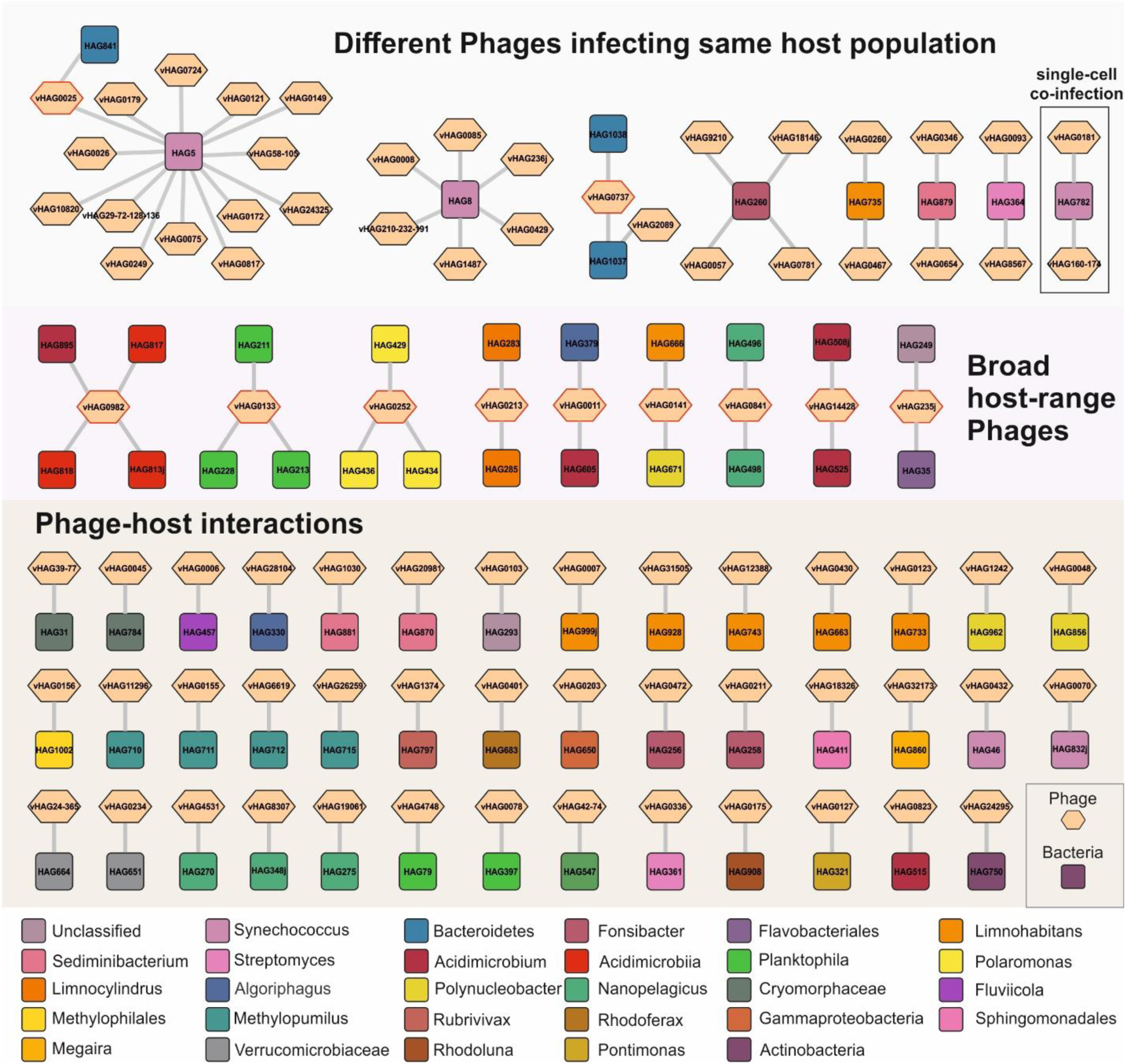
Phage-host interaction network of the AmaJul24 dataset based on Hi-C linkages. Squares and hexagons represent bacterial and viral genomes, respectively. The color of bacterial nodes indicates the taxonomy of the host. For vHAG nodes, the red outer ring denotes broad host-range phages.

### Conserved infection traits in phages targeting a shared host population

Within the linkage-derived active infection network, eight groups were identified in which two or more viruses were linked to the same host population (affiliated with *Synechococcus* (n=3), *Streptomyces*, *Limnohabitans*, *Ca.* Fonsibacter, Bacteroidetes, and *Sediminibacterium*) (Figure 2). These linkages denote an active infection pressure exerted by multiple vHAGs on a single sensitive host population (HAG) within the community, resembling a potential kill-the-winner dynamic.

Conserved genomic features (cut-off: 30% similarity over 50 amino acids) were detected among vHAGs linked to *Synechococcus* HAG5, HAG8, and HAG782, which are targeted by 14, 5, and 2 vHAGs, respectively (Supplementary Table S8). Across these groups, shared genomic regions within each group predominantly contained hypothetical proteins (*n=*12, 21, and 2, respectively); however, several proteins with known functions were also conserved (Figure 3). vHAGs linked to HAG5 shared two short tail fibers, an endolysin outer membrane protein, and a metallo-phosphoesterase. Among those linked to HAG8, three endolysins (two showing 60% similarity over 42 amino acids), a tail fiber protein, an integrase, a DNA primase, a SAM-dependent methyltransferase, and a DarB-like anti-restriction protein were shared. Between vHAG160-174 and vHAG0181, both linked to *Synechococcus* HAG782, seven conserved key regions were identified, encoding for a tail protein, a tail tube, an endolysin, two HNH endonucleases, and two hypothetical proteins.

**Figure 3.**
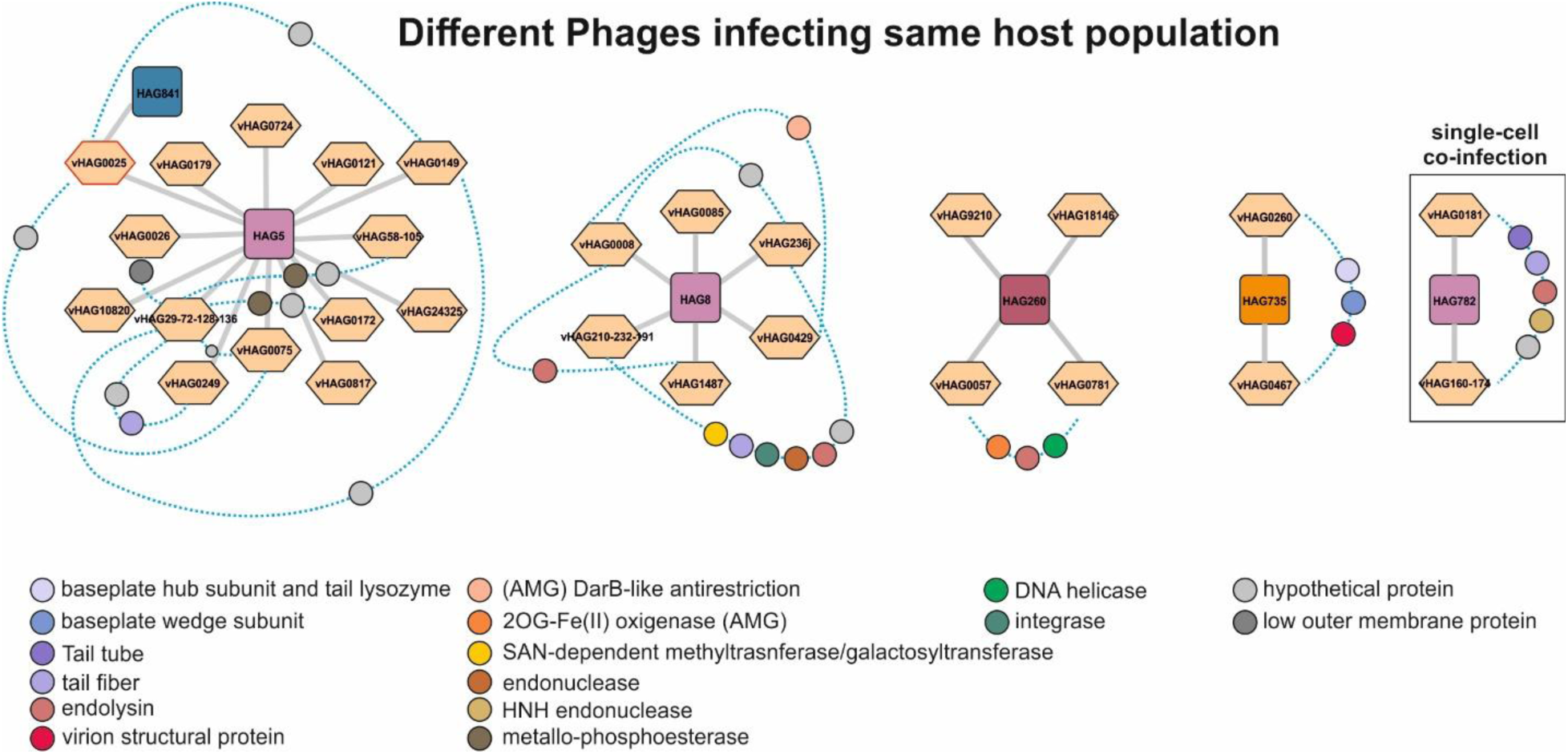
Conserved proteins across vHAGs infecting the same host HAG in the AmaJul24 dataset. Each network depicts groups of vHAGs (hexagons) infecting a shared host HAG (squares). Light blue dashed lines connect phages that share conserved protein-coding genes, which are represented as colored circles placed along each line. Circles are color-coded according to the protein function (legend below). The color-code for HAG taxonomy is the same as in Figure 2.

Similarity of tail fiber proteins among different vHAGs infecting the same host population, given the role of tail fiber as a key determinant of phage host range^43,44^, underscores the importance of maintaining efficient adsorption mechanisms leading to their specificity towards this host in the ecologically competitive freshwater environment. Likewise, the conservation of endolysins across different vHAGs linked to the same host suggests fine-tuning of the lysis strategies to the host’s cell wall architecture. These shared modules indicate that vHAGs infecting the same host population collectively shape the intensity of top-down control exerted on their host. Notably, the identification of conserved host-modulatory elements (VFDB virulence factors, HNH endonucleases, SAM-dependent methyltransferases, and DarB-like anti-restriction proteins) among these vHAGs suggests that, in addition to exerting lytic pressure, these phages potentially influence host function and genome evolution by modulating its metabolism and enabling immune evasion, potentially promoting host survival and indirectly enhancing phage persistence. Finally, the conservation of integrases and DNA primases among vHAGs associated with HAG8 suggests a potential capacity for lysogeny or chronic infection, allowing these phages to persist throughout the host population dynamics, transient blooms, or fluctuating environmental conditions. Together, these shared genomic features across distantly related vHAGs linked to *Synechococcus* populations reflect a dual pattern of conserved core infection functions and acquisition of adaptive accessory genes. The genetic profile of these vHAGs underscores the pivotal role of phages in shaping *Synechococcus* population dynamics, a key freshwater primary producer at the base of the food web.

Beyond vHAGs that retain conserved infection modules to optimize host interactions within each group, some may also represent lineages adapting to suboptimal hosts. Such adaptations could enhance their capacity for top-down control in a kill-the-winner function, thereby influencing their host-HAG population dynamics^45^. Supporting this view, evidence of episodic diversifying selection was detected in several genes from vHAGs associated with *Synechococcus* HAG5 and HAG8. In the HAG5 group, 35 genes within 25 different orthogroups encoded in eight vHAGs (vHAG0026, vHAG58-105, vHAG29-72-128-136, vHAG0172, vHAG0075, vHAG0121, vHAG0149, vHAG0249) exhibited significant signatures of diversifying selection (Supplementary Table S9). These diversifying genes encode virion structural proteins, major head and head scaffolding proteins, short tail fiber proteins, tail sheath stabilizers, enzymes involved in porphyrin biosynthesis, metallo-phosphoesterases, RusA-like Holliday junction resolvases, and several hypothetical proteins. Similarly, in the HAG8 group, a total of 20 genes distributed across 13 orthogroups (found in vHAG236j, vHAG210-232-191, vBIN0085, vBIN0008) showed also significant evidence of diversifying selection, but majority of them remain unannotated. Notably, two genes encoding tail fiber proteins in vHAG236j and vHAG210-232-191 showed particularly strong signatures of episodic diversifying selection (dN/dS ratio, ω). For vHAG236j, the maximum ω was 287.99 (*p-value=* 0.000), and for vHAG210-232-191, 778.92 (*p-value=*0.000). The detected episodic diversifying selection among phages detected as linked to *Synechococcus* populations can be a representation of the kill the winner hypothesis within the community, where these phages are quickly adapting to infect the fast-growing bacterial population or preparing for a decline in their host population through diversifying.

In the case of vHAG0260 and vHAG0467, both linked to *Limnohabitans* HAG735, shared genomic regions contained genes coding for three baseplate wedge subunits, a virion structural protein, a baseplate hub subunit, and a tail lysozyme (Supplementary Table S8). All these conserved genes were related to structural components of the virion. The presence of these conserved structural modules likely reflects neutral selection to stabilize an efficient infection apparatus, along with a conserved lysis function to ensure timely progeny release. Only three genes encoded by the vHAG0260 within three different orthogroups showed significant episodic diversifying selection. While two of these could not be annotated, one gene encoded a baseplate wedge subunit. Adaptive amino acid substitution in one copy of this subunit (max ω=2833.04, *p-value*=0.043) within the vHAG0260, which encodes baseplate wedge subunit genes clustering into three different orthogroups, reiterates the on-going arms race between the phage and its host, and potentially this phage ventures to infect new hosts for increased fitness while maintaining efficient interaction with its current host. This can be another representation of kill-the-winner where phages capitalize on the existing fast growing host population to increase their fitness while maintaining a diverse host range.

Among the four vHAGs linked to *Ca.* Fonsibacter HAG260, only vHAG675 and vHAG6781 shared genomic regions coding for a DNA helicase, two endolysins, and a 2-oxoglutarate-Fe(II) oxygenase, a versatile enzyme family involved in oxidative modifications of nucleic acids and proteins. In the context of phage infection, such oxygenase could potentially contribute to host DNA modification, counteracting bacterial defense systems, or assist in reprogramming their host’s metabolic pathways to favor viral replication^46^. None of the genes within these vHAGs showed evidence of significant diversifying selection.

### Uncovering cellular-level co-infection events through Hi-C linkages

Screening of Hi-C linkages between vHAGs associated with the same host revealed an interaction between two viral genomes linked to *Synechococcus-*HAG782, vHAG160-174 and vHAG0181 (Figure 4). According to CheckV, vHAG0181 is 100% complete with a genome size of 45.7 Kb, while vHAG160-174 is 89.7% complete and spans 84.1 kb. Hi-C analysis revealed 16 linkage events between the two vHAGs, compared to 2,142 and 1,257 intra-vHAG links for vHAG160-174 and vHAG0181, respectively (Figure 4b and c). These results strongly support the interpretation that the two vHAGs represent genetically distinct phages co-infecting the same *Synechococcus* host cell.

**Figure 4.**
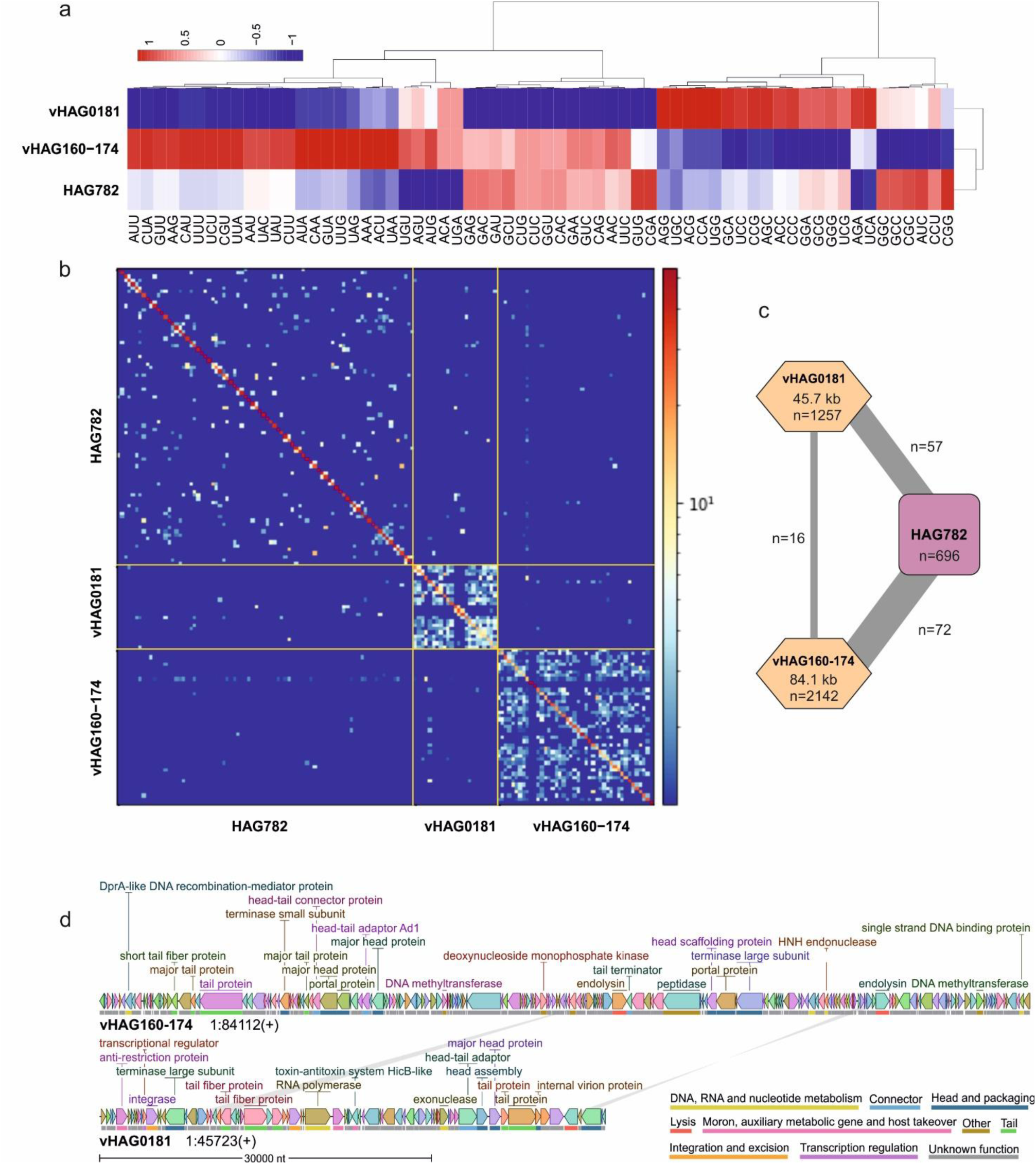
Viral co-infection event in *Synechococcus* HAG782. (a) Heatmap comparing codon usage profile of vHAG160-174, vHAG0181 and their shared host HAG782. Red indicates higher similarity, and blue indicates lower. (b) Hi-C contact map showing physical interactions between the host HAG782 and the two vHAGs. The strong signal along the diagonal corresponds to intra-genome linkages, while off-diagonal signals reflect virus–host interactions. (c) Co-infection network linking vHAG0181 and vHAG160-174 with *Synechococcus* HAG782. The thickness of the grey edges is proportional to the number of Hi-C links between bins; link counts are indicated on the edges. Number of links within the HAG and each vHAG are also written as n. (d) Genome comparison between vHAG160-174 and vHAG0181. Shared genomic regions are connected by grey bars, and genes are color-coded according to functional category (legend down right side).

These two vHAGs possess a markedly different GC-content (51.2% for vHAG160-174 *versus* 64.7% for vHAG0181), while their host *Synechococcus-*HAG782 GC-content is 59.9%. Intriguingly, these two vHAGs display almost opposite codon usage patterns (Figure 4a), but vHAG160-174 clusters more closely with the host regarding the codon usage profile, which suggests different levels of host adaptation between the two vHAGs. This pattern may indicate that vHAG0181 has either recently expanded its host range to *Synechococcus-*HAG782, or is a phage with a broader host range that is under weaker selective pressure for codon optimization. Both vHAGs show higher relative abundance in the AmaJul24 metagenome (6,043 rpkgs for vHAG160-174 and 3,913 for and vHAG0181) compared to their host *Synechococcus-*HAG782 (576 rpkgs), indicating a virus-to-host ratio greater than 1:1 and consistent with a lytic infection strategy or maybe potentially a wide host range. Nevertheless, the difference in virus-to-host ratios between the two vHAGs may reflect variation in infection dynamics, including differences in burst size, infection timing, phage-host interaction efficiency, or yet unknown phage-phage interactions. The co-occurrence of both vHAGs at high abundance, coupled with similar Hi-C linkage density, suggests they may coexist through niche differentiation^47^. Such dynamics are consistent with the “within-host coexistence model”, in which competing phages could potentially persist by balancing trade-offs in adsorption rate, latency, progeny release strategy, or burst size^48–50^. While Hi-C linkages confirm physical proximity within the same host cell and co-infection, further information regarding the infection dynamics of these vHAGs is required to disentangle their interactions.

Two small regions of high sequence similarity were detected between vHAG160-174 and vHAG0181, encoding an endolysin (54%) and a tail tube (78%), as predicted by Phold structural annotations (Figure 4d, Supplementary Table S8). Sharing a similar endolysin and tail tube may reflect conservation or horizontal gene transfer of a lysis and identification module, both likely optimized for the *Synechococcus* host cell wall. Moreover, these conserved functional proteins could facilitate synchronized or complementary lysis timing between the two phages, potentially indicating cooperative or competitive interactions during co-infection. The presence of a shared endolysin gene within otherwise divergent genomic backgrounds suggests that modular recombination may serve as a mechanism of adaptive gene exchange among these two vHAGs and freshwater phages at large, consistent with patterns reported for T4-like cyanophages^51^ and the extensive gene flow observed in aquatic viral communities^4,52,53^. As mentioned above, other shared regions were also detected between these two phages, encoding for tail protein, HNH endonucleases, and hypothetical proteins.

Among all genes of these two phages, only a gene encoding a deoxynucleoside monophosphate kinase (max ω= 2.51905E+18, *p-value*=0.033) and a gene encoding a hypothetical protein (max ω=6430.62, *p-value*=0.001), both present in vHAG160-174, were detected to be under significant diversifying selection (Supplementary Table S9). This hypothetical protein belongs to an orthogroup together with another gene from vHAG160-174 (not under significant diversifying selection) and two additional genes from vHAG210-232-191 (under significant diversifying selection, max ω=15114.87, *p-value*=0.014) and vHAG236j (not under significant diversifying selection), both linked to *Synechococcus* HAG8. No gene within this orthogroup could be functionally annotated; however, the fact that it contains two genes encoding proteins with an amino acid identity of 87%, and that only one of them shows evidence of adaptive amino acid substitutions in vHAG160-174, suggests a yet unknown but important evolutionary role for this hypothetical protein, most likely related to its interactions with its host.

Similarly, the vHAG160-174 encodes two genes that belong to the same orthogroup annotated as deoxynucleoside monophosphate kinases (89% amino acid identity). However, only one of them undergoes significant diversifying selection. Phage-encoded deoxynucleoside monophosphate kinases are common and have been reported in jumbo phages^54,55^, and also marine cyanophages^56^. In this orthogroup, homologous genes were detected in two jumbo phages (vHAG0252, 260.3 kb; vHAG235J, 413.6 kb) as well as in two copies in the cyanophage vHAG160-174. This enzyme is involved in phage replication by providing nucleotide precursors or enabling phage genome replication once the host metabolism is shut down during the infection. The presence of two copies of this gene in a phage infecting a single host cell co-infected by two phages, one undergoing significant diversifying selection, highlights potential competition between these phages to access the nucleotide pool required for replicating their genomes.

### Genomic Signatures of Broad-Host-Range Phages in Freshwater Viral Networks

Hi-C linkages detected eleven vHAGs associated with more than one host, indicating the prevalence of broad host range or generalist phages actively infecting diverse freshwater lineages. The host range of these phages extended from populations within the same genus (*Ca.* Nanopelagicus, *Ca.* Limnocylindrus, *Polaromonas*, *Ca.* Planktophila), to members of the same family (Burkholderiaceae), class (Acidimicrobiia), phylum (Bacteroidota), and even across distinct bacterial phyla, including Bacteroidota and Cyanobacteriota, Bacteroidota and Actinomycetota, as well as Bacteroidota and an unclassified lineage (Figure 2)^25,32^. Among these, vHAG235j was predicted to be an integrase containing phage; nevertheless, it showed a virus-host ratio greater than one compared with both of its target hosts. This can be an outcome of its broad host range or a lytic infection in one or both hosts.

Broad-host-range viruses breaching into hosts taxonomically far apart are increasingly being described in literature and isolated in the laboratory^22,32,57,58^. Broad-host-range phenotype has also been shown to be driven by chimeric infectious particles generated by tail hijacking by capsid-forming “Phage-Inducible Chromosomal Islands” or PICIs^29^. Moreover, a recent study using Hi-C proximity ligation on deep-sea microbial mats also reported viruses interacting with hosts from different classes or even across domains^57–59^. In freshwater ecosystems, where host densities are often low, such broad-host-range interactions may reflect ongoing viral adaptations to suboptimal hosts as a strategy to sustain their own viral population^45^.

Phage-exerted top-down pressure is known to shape genomic heterogeneity within host populations^60^. Broad-host-range and generalist phages may impose diffuse selective pressures across multiple host genotypes, potentially promoting higher intra-population diversity. In contrast, infection by highly specific viral populations might lead to stronger population bottlenecks, thereby reducing microdiversity^61^. However, comparison of the microdiversity profiles of HAGs infected by generalist vHAGs and those infected by specialist vHAGs revealed no significant differences between the two groups (Mann-Whitney U test, W = 125650, *p-value*=0.8099) (Supplementary Figure S6).

Analysis of the dN/dS ratios for genes from these broad-host-range phages revealed evidence of significant diversifying selection in 29 genes belonging to 25 different orthogroups encoded by vHAG235j, vHAG0011, vHAG0252, vHAG0133, and vHAG0982. While the majority of these proteins remain unannotated (n=20), several genes with annotated functions were also identified, including a baseplate hub subunit, a tail lysozyme, a baseplate wedge tail fiber protein connector, an endolysin, a major head protein, a major tail protein, a portal protein, both terminase subunits, and a virion structural protein. Interestingly, in all these cases, the same phage also encoded another copy of the same gene within the same orthogroup with no detectable amino acid level adaptation (Supplementary Table S9). This is specifically important for genes encoding key structural components, which must preserve infection efficiency while simultaneously enabling adaptive diversification through gene duplication, a known mechanism that seems to be prevalent among freshwater phages.

### Temporal dynamics of linked phage-host populations

Within the AmaJul24 metagenome, pairwise comparisons of the abundance of host HAGs infected by generalist phages (*n=*22), by multiple distinct viral populations (*n=*36), or involved in a single virus-host interaction (*n=*42), showed no significant differences (Wilcoxon signed-rank test with Bonferroni correction, *p-value* = 1 for all comparisons). In contrast, pairwise comparison of vHAG abundances indicated that vHAGs linked to a single host were significantly less abundant than those infecting a sensitive host population targeted by multiple phages (Wilcoxon signed-rank test, *p-value* = 0.0082), and showed lower abundances compared to generalist vHAGs (albeit not significant, Wilcoxon signed-rank test, *p-value* = 0.15). Phages targeting sensitive hosts and generalist phages showed a similar abundance pattern (Wilcoxon signed-rank test, *p-value* = 0.8). The virus-to-host abundance ratios (VPH) of the generalist phages were significantly higher than those infecting sensitive hosts (Wilcoxon signed-rank test, *p-value* = 0.00078), whereas differences in the VPH ratio of generalist phages and single-interaction viruses were not significant (*p-value*=0.15 and 0.21, respectively). These results suggest that killing the winner host population can lead to significant higher abundances among the vHAGs infecting these sensitive host populations compared to the single virus-host interactions, even though these sensitive hosts were not significantly more abundant than hosts infected by a single virus and did not lead to higher VPH ratios. Thus, our results reiterate that the factor defining the intense phage control on these winner populations is attributed to their replication rate and not reflected in their final abundance in the community^62^.

To assess the temporal stability of phages and their co-evolutionary dynamics with their host, vHAGs were classified as “stable” if detected in 12 available metagenomes from Amadorio collected on 4 timepoints over the span of 13 years (including 25 vHAGs infecting sensitive host, 6 generalist vHAGs, and 14 vHAGs associated with single hosts), and as “transient” if present in a subset of these metagenomes (including 8 vHAGs infecting sensitive host, 5 generalist vHAGs, and 27 vHAGs associated with single hosts) (Supplementary Table S10). Temporal abundance of linked hosts and phages across different metagenomes sequenced from the Amadorio reservoir shows a stronger correlation for sensitive and broad host range phages and their host, whereas one-to-one infections had a wide range of correlation values (Supplementary Figure S7). Stable vHAGs were consistently more abundant than transient vHAGs (Wilcoxon test = 205298, p-value < 2.2e-16), the more transient phages may thrive under episodic conditions or within competitive niches structured by phage–phage and phage–host coevolution, or by infecting suboptimal hosts.

In terms of linkages, 46 phage-host relationships were formed by transient vHAGs and stable HAGs, two by transient vHAGs and transient HAGs, 49 by stable vHAGs and stable HAGs, and three by stable vHAGs and transient HAGs (Supplementary Table S11). Two of the transient HAGs were infected by a stable broad host range phage, and another one was infected by a single stable vHAG. These observations suggest a potentially broader host range for these phages beyond detected linked HAGs.

For the two vHAGs co-infecting *Synechococcus-*HAG782 at the cellular level, all three counterparts are detected in all Amadorio metagenomes, suggesting continuous interaction. However, vHAG0181 shows a stronger abundance correlation with its host (0.45) compared to vHAG160-174 (-0.09). These results from snapshot view provided by the available metagenomes suggest a temporal variation in their interactions, possibly impacted by their differing burst sizes, infection strategies, or replication rate of the host lineage in different seasons and timepoints (Figure 5). On a global scale though the abundance correlations of the host with vHAG0181 and vHAG160-174 are higher (0.81 and 0.80, respectively), which signify their strong three-party interaction in different lakes (Supplementary Figure S8).

**Figure 5-.**
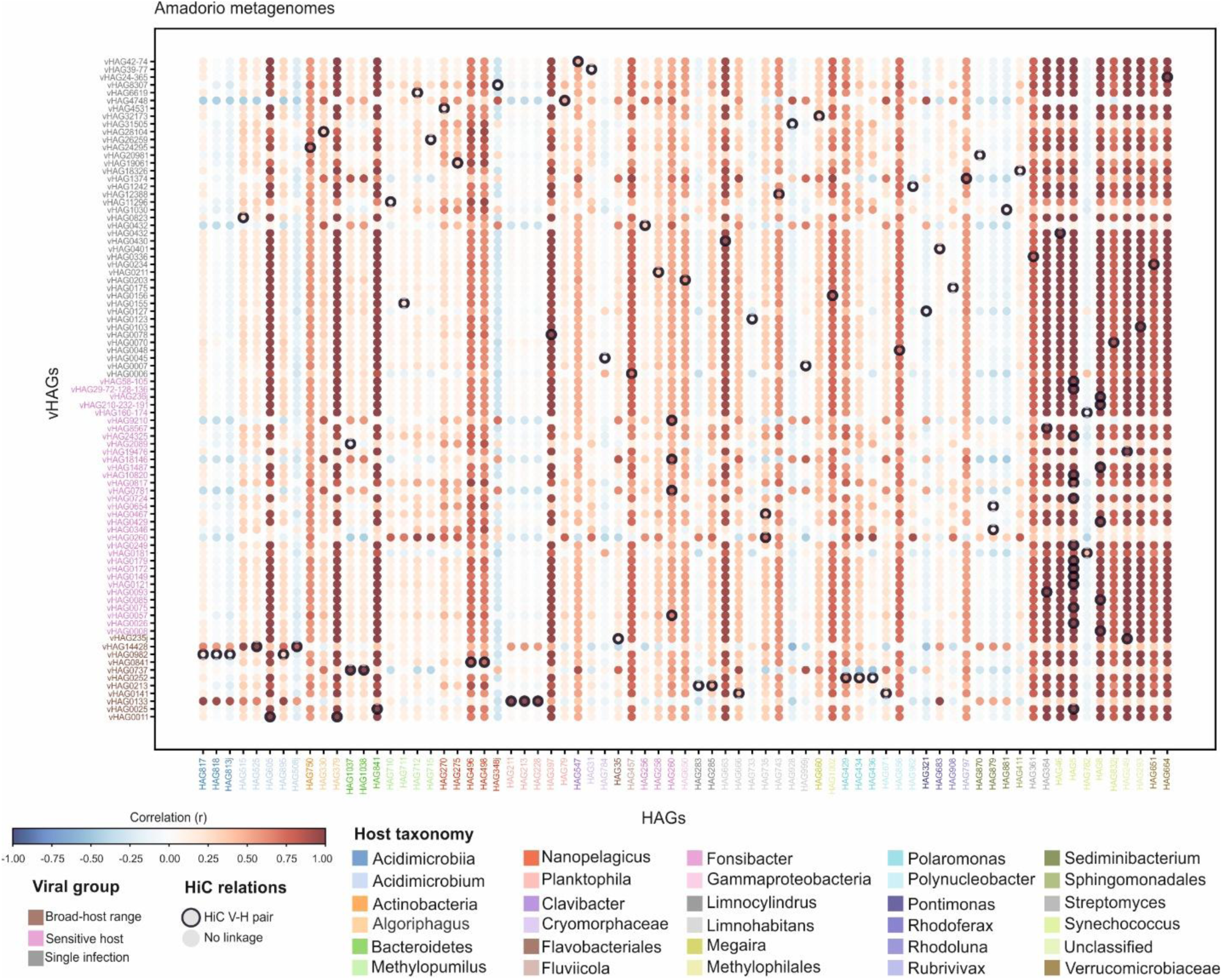
HAG and vHAG abundance correlation. Correlation of linked HAGs and vHAGs abundances across 12 available metagenomes from Amadorio Reservoir from 2012 to 2024, sampled at different depths (0, 5, 10, and 20 m).

### Conclusion

Over the last decade, the discovery of bacterial defense systems and corresponding phage counter-defense strategies has advanced our understanding of phage–bacteria co-evolution. Despite these advances, how such co-evolutionary processes are embedded within the ecological dynamics of natural microbial communities remains insufficiently resolved. A major limitation has been the difficulty of identifying active phage–bacteria infections in situ and assessing how these interactions vary over time. MetaHi-C–based linkage analysis provides an approach to address this limitation by enabling the detection of phage–host active interactions directly within complex communities which is then enabling us to link these ecological dynamics to underlying evolutionary processes. Applied to freshwater systems in this study, this framework provides a direct view of the intertwined ecological (phage host range, killing-the-winner, cellular level co-infection, abundance correlations) and co-evolutionary (conserved infection modules versus diversifying selection in host interaction genes) processes that shape freshwater population diversity and dynamics and establishes a basis for extending similar analyses to other natural microbial communities.

## MATERIAL AND METHODS

### Sample collection

Approximately 30 L of water were collected on July 25, 2024, from the Amadorio freshwater reservoir in Alicante, Spain (38° 32′ 1.653″ N, 0° 15′ 53.252″ W). The water sample was taken at a depth of 5 m and prefiltered through 20 µm pore size filters. From the resulting filtrate, 100 mL was used to prepare two Hi-C libraries. The remaining volume was filtered through 0.22 µm pore size polycarbonate filters (Millipore) to collect microbial biomass for DNA extraction and sequencing. All filters were stored at –80°C until further processing.

### DNA extraction, shotgun library generation, and sequencing

Metagenomic DNA was extracted from the 0.22 µm filters (containing the biomass of 5 L of the freshwater) with DNeasy Blood and Tissue kit (Qiagen, Hilden, Germany). Genomic DNA was processed with the Nextera XT Library Preparation kit and sequenced on an Illumina NovaSeq6000 (PE150), yielding a reference metagenome of 89.4 Gb (596 million paired-end 150 bp reads). This metagenomic sample is hereafter referred to as AmaJul24 (Supplementary Table S12).

### Hi-C Library Preparation and Sequencing

Cells from 50 mL of the collected samples were concentrated into a single pellet by centrifugation at 75,600 x g for 40 min. Two technical replicates were generated from the same sample. Each pellet was processed using the ProxiMeta Hi-C Microbiome v.4.0 Kit (Phase Genomics, Seattle, WA). Following the manufacturer’s protocol, crosslinked DNA was digested with the restriction enzymes *Sau3AI* and *MluCI*, enabling the formation of chimeric DNA molecules via proximity-based ligation with biotin-tagged nucleotides. These chimeric molecules, representing physically proximal genomic regions, were selectively captured using streptavidin-coated beads and further processed using the ProxiMeta library preparation reagents. Sequencing was performed on an Illumina NovaSeq 250PE platform, generating 34.1 Gb of data (FW1, 73 million reads; FW2, 63 million reads). After quality control, both datasets were merged into a single library, AmaJul24-HiC (136 million reads, Supplementary Table S12).

### Metagenomes and Hi-C data processing

Quality control of both AmaJul24 and AmaJul24-HiC metagenomes was performed using *bbduk* from the BBTools suite (v.37.25)^63^. Quality-controlled metagenomic reads from AmaJul24 were assembled into contigs using MEGAHIT (v.1.2.9)^64^, with the following settings: ‘-min-contig-len 1000 -k-min 21 -k-max 141 -k-step 12 -merge-level 20, 0.95’. The assembly resulted in 380,079 contigs. The Megahit_toolkit (v.1.2.9) was used to obtain the fastg assembly graph from the K141 k-mer intermediate. Assembly graphs were visualized using Bandage (v.0.8.1)^65^. Viral contigs larger than 1 kb were identified from the assembly using both Vibrant (v.1.2.0)^66^ and geNomad (v.1.7.1)^67^ with default parameters. AmaJul24-HiC reads were aligned to the assembled contigs using BWA-MEM (v.0.7.17)^68^, with the ‘-5SP’ flag. Reads that were unmapped or flagged as secondary, supplementary, or of poor quality (mapping score or aligned length < 30), were discarded. Raw Hi-C contact maps were generated by counting the number of Hi-C read pairs mapping to each contig pair.

### Reconstructing HiC-Assembled Genomes (HAGs and vHAGs)

Hi-C reads were used to improve the quality of the HiC-Assembled Genomes. Contigs from the AmaJul24 assembly larger than 1 kb were binned using MetaCC (v.1.0.0)^69^, with a minimum bin length of 30 kb and specifying the restriction enzymes used in the AmaJul_HiC preparation. Bin quality was assessed with CheckM (v.1.2.2)^70^ with the *lineage_wf* workflow, and only HAGs with ≤5% contamination^33^ were retained for further analysis.

ViralCC (v.1.0.0) was employed to obtain Viral HiC-Assembled Genomes (vHAGs) by integrating the Hi-C interaction graph with a host proximity graph for viral contigs. Only viral contigs identified by both Vibrant (v.1.2.0) and geNomad (v.1.7.1) were used as input for ViralCC, applying a minimum contig length of 1 kb. CheckV (v.1.0.1)^34^ was used to estimate the quality of each vHAG. Prokaryotic HAGs were clustered into genomospecies using dRep (v.3.5.0)^71^ with the “compare” option, default parameters, and a 95% ANI threshold. vHAGs were then concatenated and clustered into species-level vOTUs (following a 95% ANI and 85% alignment fraction of the shorter sequence^72^).

### Optimizing the HiC-Assembled Genomes

The binning software MetaCC initially separated contigs from the same HAG/vHAG into different bins due to variation in linkage density across genomic regions. As previously seen^37^, prokaryotic DNA presents a well-established 3D conformation in CIDs. Consequently, MetaCC can assign contigs from the same HAGs/vHAGS to separate bins corresponding to these CIDs, reflecting the underlying 3D genome conformation. This folding of DNA results in genomic regions with variable linkage densities that MetaCC may incorrectly interpret as belonging to different genomes (or bin them individually), leading to artificial over-fragmentation. To address this, all HAGs were systematically compared to identify linkages between them. To evaluate linkage density within and between HAGs pairs, we developed the Nadal Index (NI), which considers both intra-HAG and inter-HAG linkage densities. It was assumed that, if the intra- and inter-HAG linkage densities were not equal, the inter-HAG density would be closer to the intra-HAG density.

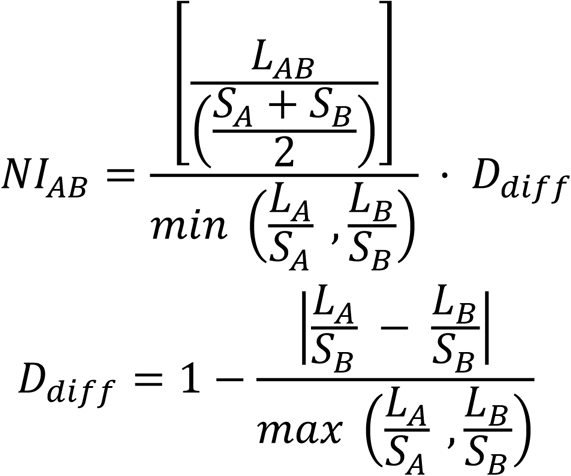

To enable comparisons of intra-HAG linkage densities, values were normalized. The resulting index ranges from 0 to 1 (though it can exceed 1). Values approaching t 1 indicate that both HAGs are likely derived from the same genome, whereas values closer to 0 indicate that they belong to different genomes. To establish a threshold, the distribution of index values was analyzed using two statistical approaches: *k-*means clustering and Otsu’s method. *K-*means produced a threshold of 0.309, while Otsu’s method yielded 0.304. Based on these results, a threshold of 0.31 was selected to define whether two HAGs belong to the same genome. Furthermore, contact maps were generated at different threshold values to validate this cut-off. It was confirmed that values below 0.31 consistently corresponded to unrelated HAGs, whereas values above this threshold were associated with HAG pairs belonging to the same genome (Supplementary Figure S9). The same method was applied to vHAGs.

To evaluate the impact of CIDs and linkage density on binning efficiency, an additional Hi-C experiment was performed using a pure culture infection of *E. coli* K12 (U00096.3) and its bacteriophage Lambda. The *E. coli* K12 reference genome was randomly fragmented in 602 fragments (N50 = 10,573 pb; longest = 36,846 pb), which were then subjected to MetaCC binning using the same parameters as previously applied. This procedure split the *E. coli* genome into five separate bins. Comparison with the *E. coli* Hi-C contact map showed that regions with higher linkage density correspond to areas where bin splits occurred, highlighting the influence of CIDs on binning (Supplementary Figure S2). These results indicate that post-binning evaluation is required to merge HAGs/vHAGs originating from the same genome, both in metagenomic datasets and in genomes derived from pure cultures (Supplementary Table S1).

### Phage-host interactions analysis using Hi–C data

Viral copies per host cell (VPH) values were estimated by combining the number of Hi-C links between a specific phage-host pair (L), normalized by the total Hi-C links that the virus has with all potential hosts (L(v)), and the ratio between phage abundance (V) and host abundance (H). To reduce false positives, phage-host pairs identified through Hi-C reads were subjected to different rounds of filtering.

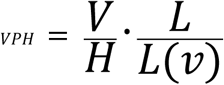

These criteria required a minimum of two Hi-C reads linking the phage and the host Hi-C assembled genome (HAG), a phage-host connectivity ratio (R’) of at least 0.1, and a minimum of 10 intra-HAG Hi-C links. The phage-host connectivity ratio (R’) was determined by comparing the Hi-C connectivity density (Hi-C link count per kilobase squared of genome) of the phage-host pair (D_VH_) with that of the HAG itself (D_H_), with both values normalized by the estimated VPH.

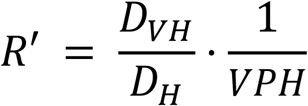

In addition, a VPH threshold derived from a receiver operating characteristic (ROC) curve was applied to maximize the detection of phages with at least one host while minimizing false phage-host associations. An initial threshold of 0.01 was determined; however, to ensure higher confidence in the resulting interactions, a final cut-off value of 0.1 was selected.

### Visualizing phage-host interaction networks

Phage-Host interaction networks were visualized using Cytoscape (v.3.9.1)^73^.

### Alternative host assignment methods for vHAGs

iPHoP (v.1.3.0) was run with default parameters and the “Aug_2023_pub_rw” database to predict the putative vHAG hosts^22^. CRISPR spacers from HAGs were extracted using MinCED (v.0.4.2)^74^ with default parameters and compared to vHAGs sequences to identify potential virus-host associations. The presence of MITEs was assessed using MITE Tracker (v.1.0.0)^75^ with default parameters, comparing viral and prokaryotic sequences at 95% nucleotide identity and 100% coverage^23^.

### Hi-C contact map generation

Hi-C contact maps were generated using the HiCExplorer (v.3.6) pipeline^76^. Read alignment was performed with BWA (v. 0.7.18-r1243-dirty) using a minimum mapping quality threshold of 30. Hi-C reads were mapped to reference genomes concatenated into a single multi-fasta file for each genome group. Contact maps were binned at 2 kb resolution for both vHAGs and HAGs, normalized to values between -1.5 and 5, and subsequently log-transformed.

### Abundance calculation via metagenome read recruitment

Fragment recruitment was carried out using Bbmap (v.38.18) to align metagenomic reads against the reference genome sequences. Recruited reads were normalized as reads per kilobase of the genome per gigabase of metagenome (rpkg). Supplementary Table S13 lists the metagenome accessions used in this study.

### Alignment of similar regions

To compare similar features between vHAGs associated with the same HAG, *tblastx* comparisons were used with relaxed parameters of >30% similarity and a minimum alignment length of 50 amino acids. For comparisons between vHAGs and viral sequences from the JGI IMG/VR database (v.4.1)^77^, protein clustering was performed with MMseqs2^78^ using the LoVis4u package (v.0.0.13)^79^.

### Microdiversity analyses

Microdiversity was analyzed using InStrain (v.1.9.0)^80^ with the “profile” pipeline. Metagenomic reads were aligned to the reference genomes using BBmap (v.38.18) to obtain pN/pS values for each gene. To obtain dN/dS values for each gene, OrthoFinder (v.3.1.0) was used to cluster all proteins into orthogroups. MAFFT (v.7.525) was then used to generate multiple sequence alignments for each orthogroup, and PAL2NAL (v.14)^81^ converted the protein alignments into codon alignments. IQ-TREE (v.3.0.1)^82^ was used to analyze the codon alignments for each orthogroup with the Goldman-Yang model^83^. Finally, HyPhy (v.2.5.75)^84^ was applied to perform selection analysis. First, the Branch-Site Unrestricted Statistical Test for Episodic Diversification (BUSTED) model^85^ was used to identify orthogroups under episodic diversifying selection. For those detected with positive selection, the adaptative Branch-Site Random Effects Likelihood model (aBSREL)^86^ was then applied to pinpoint specific genes within orthogroups undergoing diversifying selection.

### GC content and Codon usage calculation

GC-content from the bacterial and viral genomes was calculated using SeqTk v.1.3-r106 (https://github.com/lh3/seqtk.git). Codon usage was calculated using an *in-house* developed script.

### Statistical analysis

Statistical analyses were conducted in R v.4.2.1 (Team RC. R: A language and environment for statistical computing. Vienna: R Foundation for Statistical Computing; 2019) and RStudio v. 2022.07.0 (Team R. RStudio: integrated development environment for R. Boston: RStudio, PBC; 2022). Kruskal-Wallis Rank Sum Test and Pairwise Wilcoxon Rank Sum Tests were calculated using kruskal.test and pairwise.wilcox.test functions, respectively, of the stats R package (v. 3.6.2) for numeric variables of paired samples at 0.95 confidence level.

## Data availability

All the sequences obtained in this work, the AmaJul24 and the two metaHi-C libraries associated (AmaJul24_HiC_1 and AmaJul24_HiC_2), have been deposited in the NCBI BioProject PRJNA1273184 (biosamples SAMN48938357, SAMN48928570; and SAMN48938196, respectively). The linked HAGs and vHAGs fasta sequences are available for download from the Zenodo repository, 10.5281/zenodo.15856862.

## Funding

This research was supported by the “VIRHOS” project, Ref. CIPROM/2021/006 (PROMETEO 2022, Generalitat Valenciana). Maliheh Mehrshad was supported by ERC grant (Grant agreement No. 101117028 — MULTIPHAGE — ERC-2023-STG to Maliheh Mehrshad). Funded by the European Union. Views and opinions expressed are however those of the author(s) only and do not necessarily reflect those of the European Union or the European Research Council Executive Agency. Neither the European Union nor the granting authority can be held responsible for them

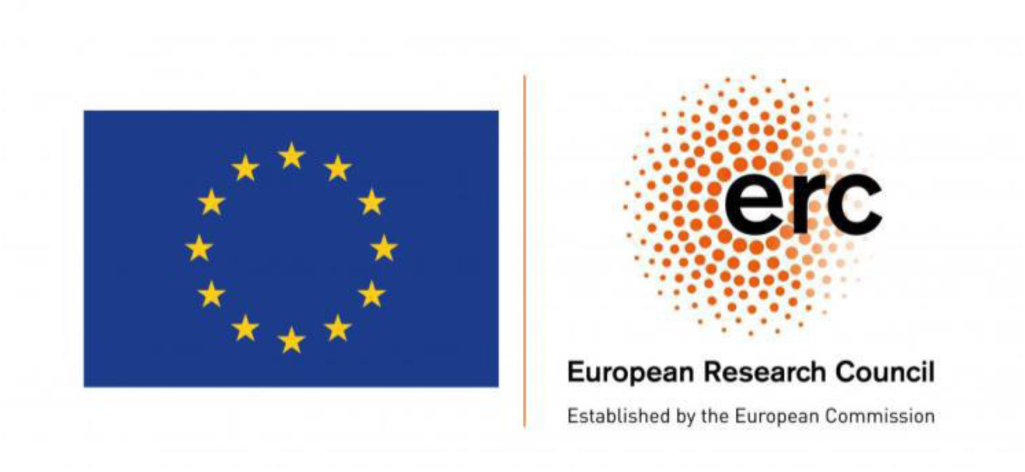

## Author contribution

FNM, ABMC, and MM designed the study. FNM and ABMC collected and possessed the samples. FNM, ABMC, and MM performed bioinformatic analysis. MM and ABMC interpreted the results and wrote the manuscript. FNM, ABMC, and MM read and approved the final version of the manuscript.

## Acknowledgements.

We thank Josefa Antón for procuring funding and the critical reading of the manuscript.

## SUPPLEMENTARY FIGURES

**Supplementary Figure S1.**
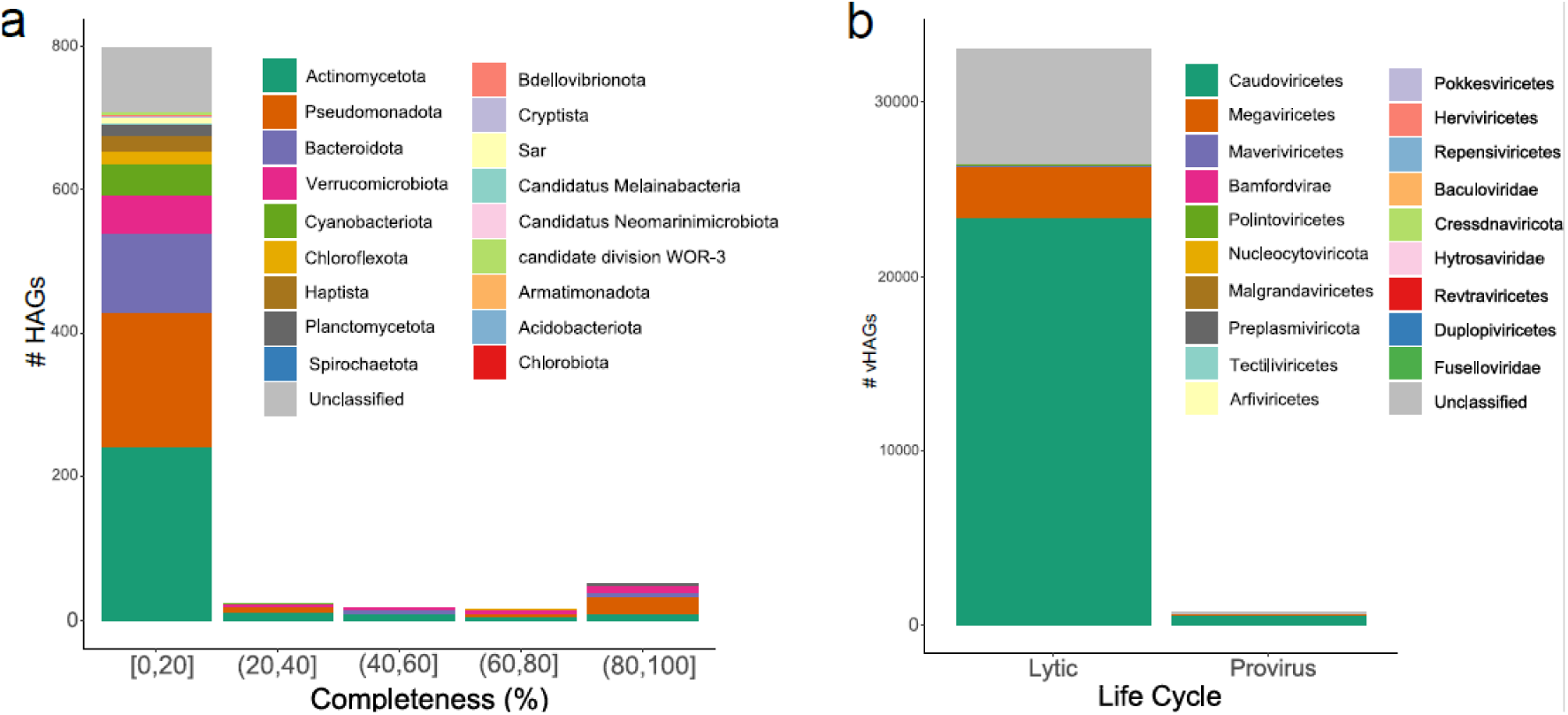
Taxonomic classification of HAGs and vHAGs in the AmaJul24 dataset. (a) Phylum-level classification of HAGs across different completeness ranges. (b) Classification of vHAGs based on predicted life cycle.

**Supplementary Figure S2.**
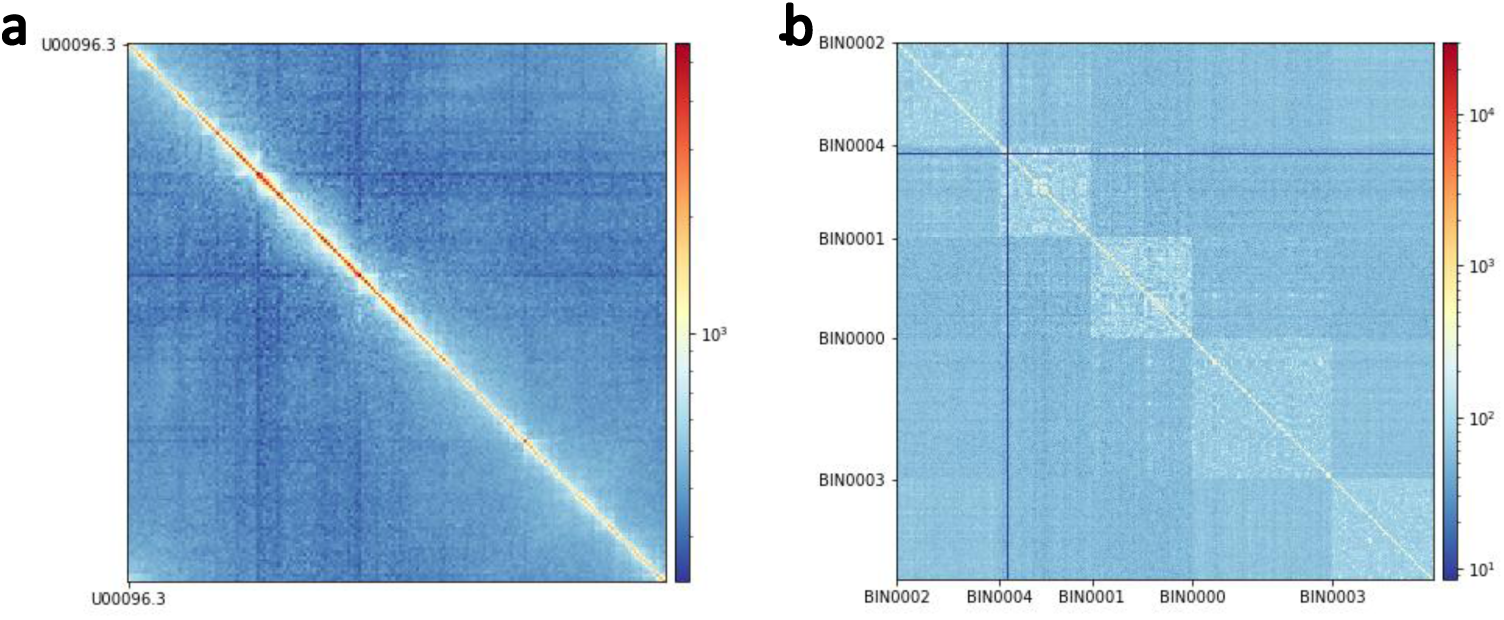
Hi-C contact maps of *Escherichia coli* K12. (U00096.3) obtained with the MetaCC program. (a) Map obtained from the complete genome. (b) Map generated after random fragmentation into 602 pieces (N50 = 10,573 pb; longest = 36,846 pb). The five resulting bins were ordered by genomic position and plotted. Regions with higher linkage density correspond to areas where bin splits occurred.

**Supplementary Figure S3.**
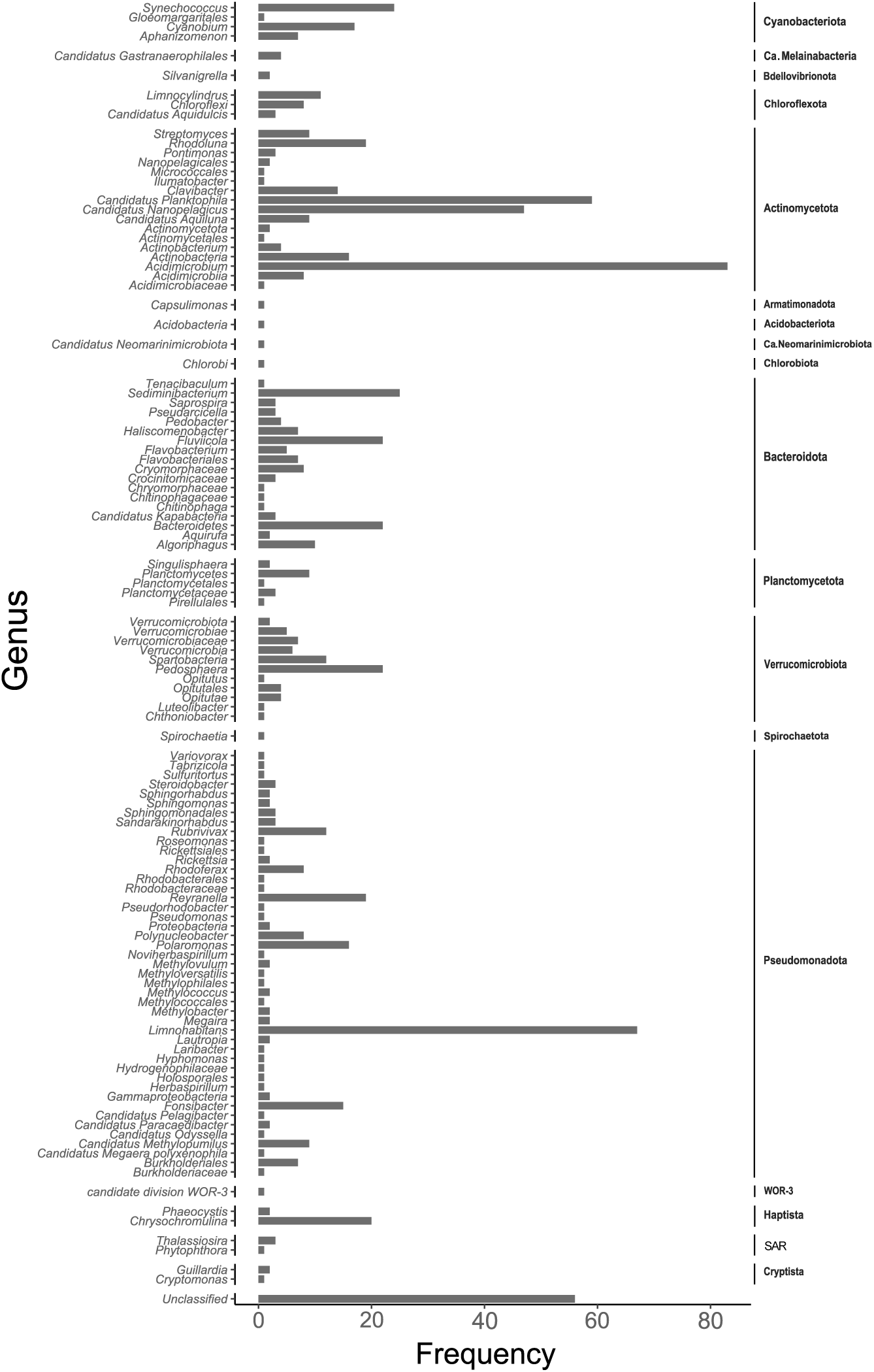
Taxonomic composition of Hi-C assembled genomes (HAGs) form the Amadorio freshwater reservoir. Taxonomic distribution of HAGs reconstructed from AmaJul24 metagenome (Alicante, Spain 2024). HAGs are grouped by phyla and genus.

**Supplementary Figure S4.**
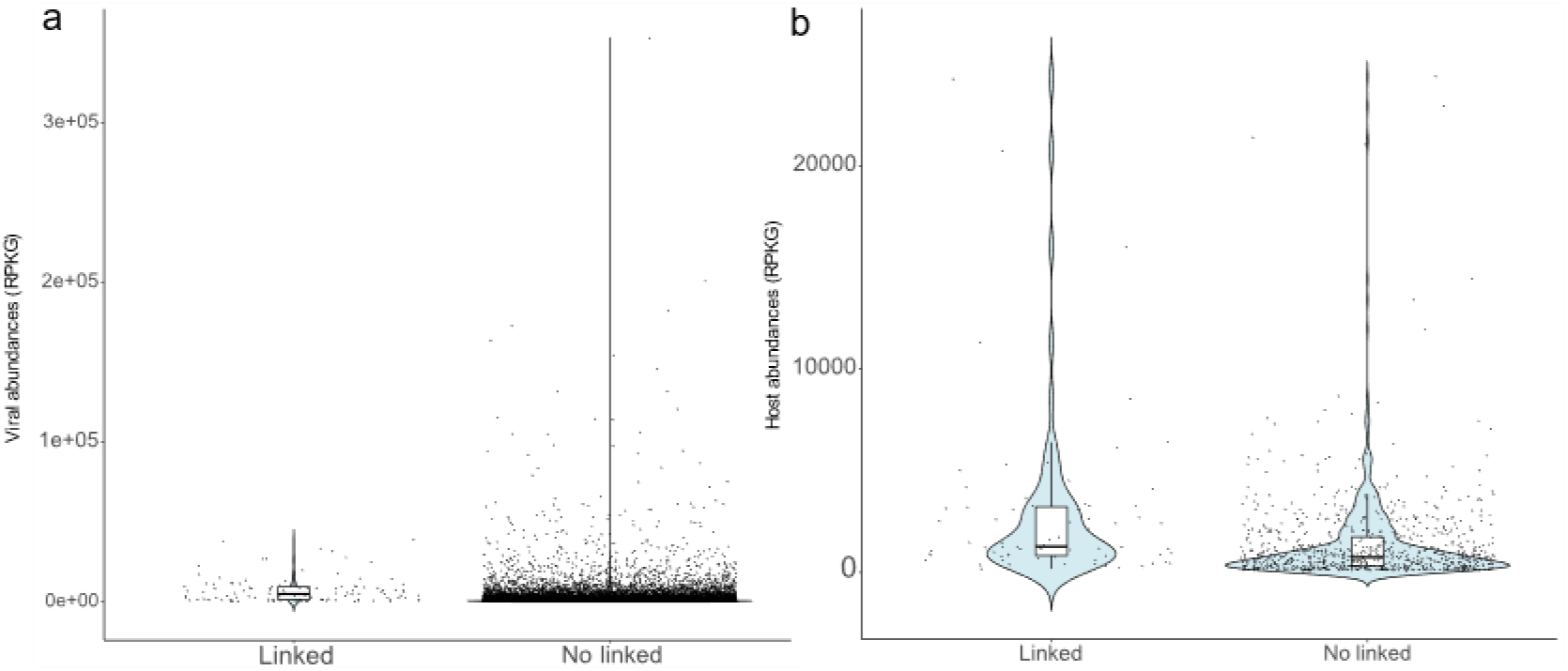
Comparison of the abundances of linked and unlinked vHAGs (a) and HAGs (b).

**Supplementary Figure S5.**
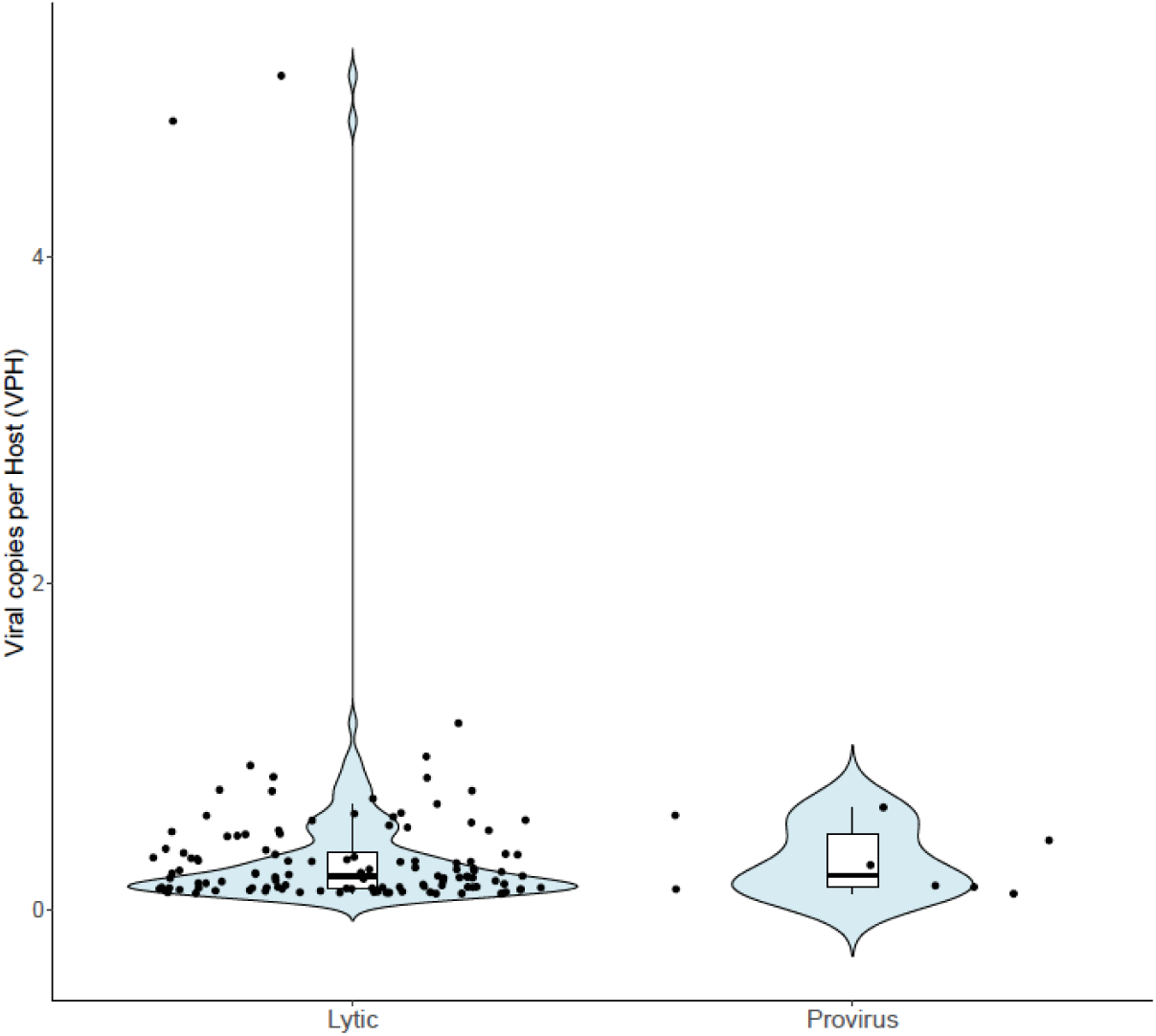
Comparison of viral copies per host (VPH) between predicted lytic and proviral vHAGs linked to HAGs. Violin plots showing the distribution of VPH values across viral populations from AmaJul24 and AmaJul24-HiC. Only vHAGs with Hi-C linkages to host HAGs were included in the analysis. The width of each violin reflects data density at different VPH values, and the central line indicates the median.

**Supplementary Figure S6.**
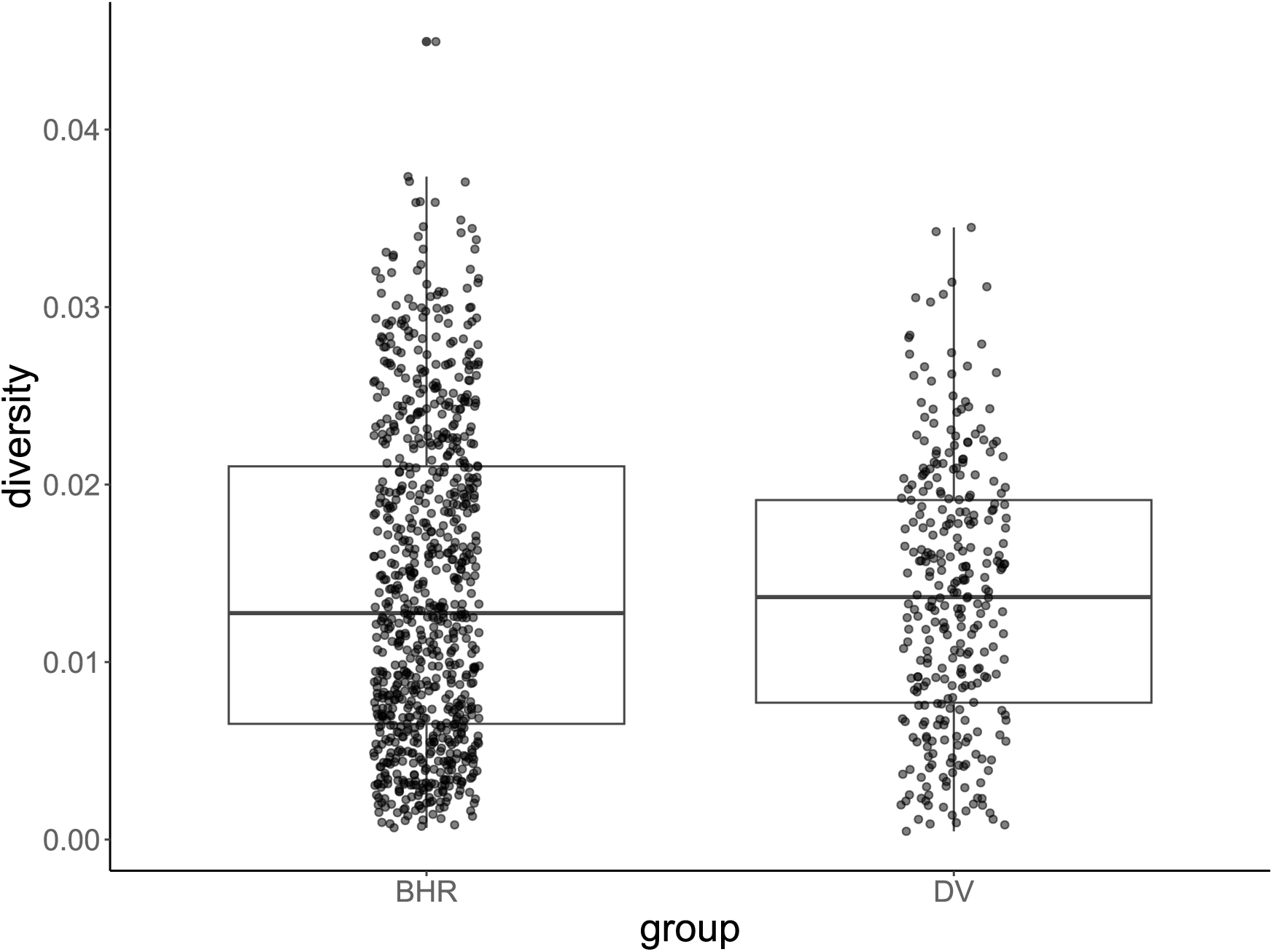
Microdiversity of vHAGs. Boxplot and jitter plot comparison between microdiversity values extracted from InStrain for broad host range vHAGs and vHAGs infecting sensitive hosts (DV).

**Supplementary Figure S7.**
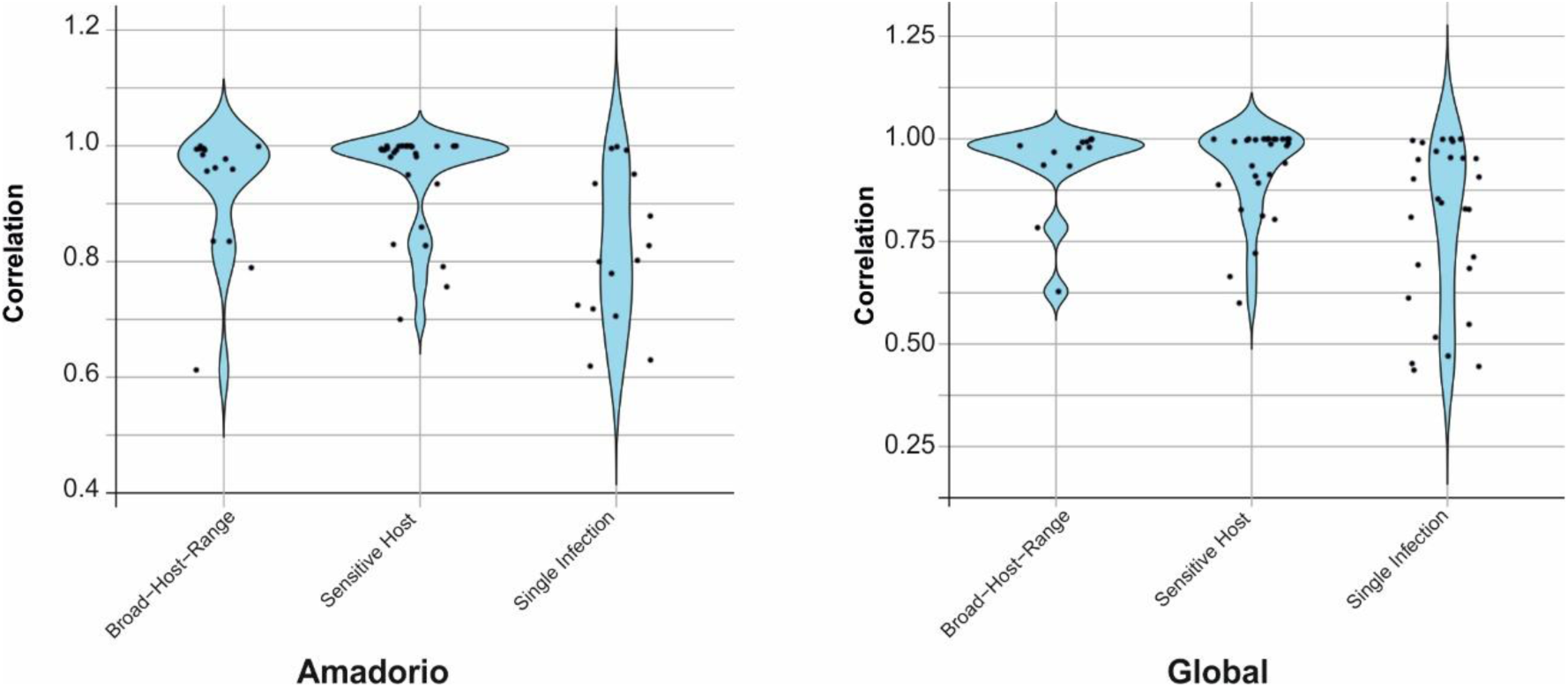
HAG and vHAG abundance correlation. Correlation comparison of linked HAGs and vHAGs abundances across 12 available metagenomes from Amadorio Reservoir from 2012 to 2024, sampled at different depths (0, 5, 10, and 20 m) (left) and 20 metagenomes across metagenomic datasets from lakes worldwide (right).

**Supplementary Figure S8.**
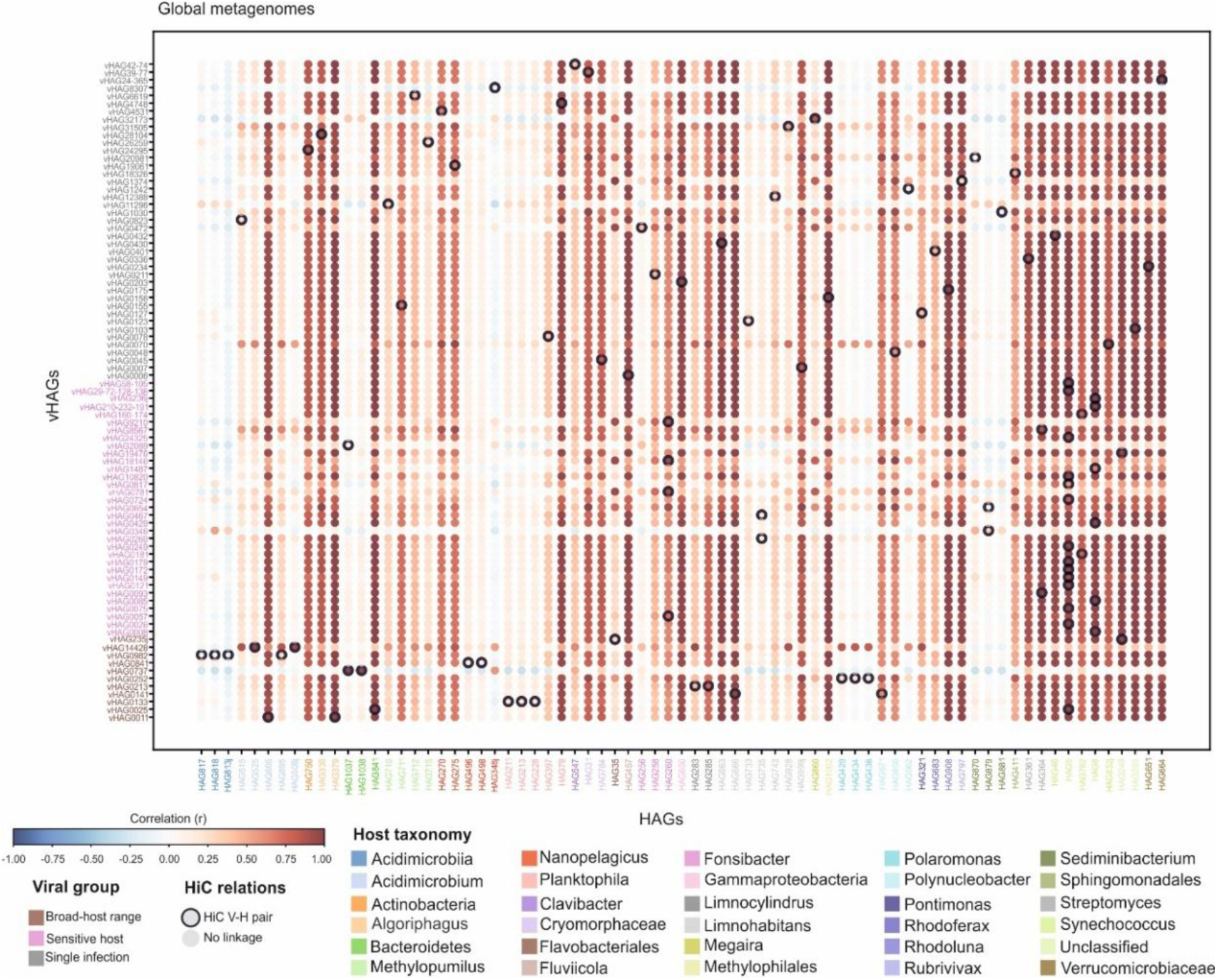
HAG and vHAG abundance correlation at global level. Correlation of linked HAGs and vHAGs abundances across 20 metagenomic datasets from lakes worldwide.

**Supplementary Figure S9.**
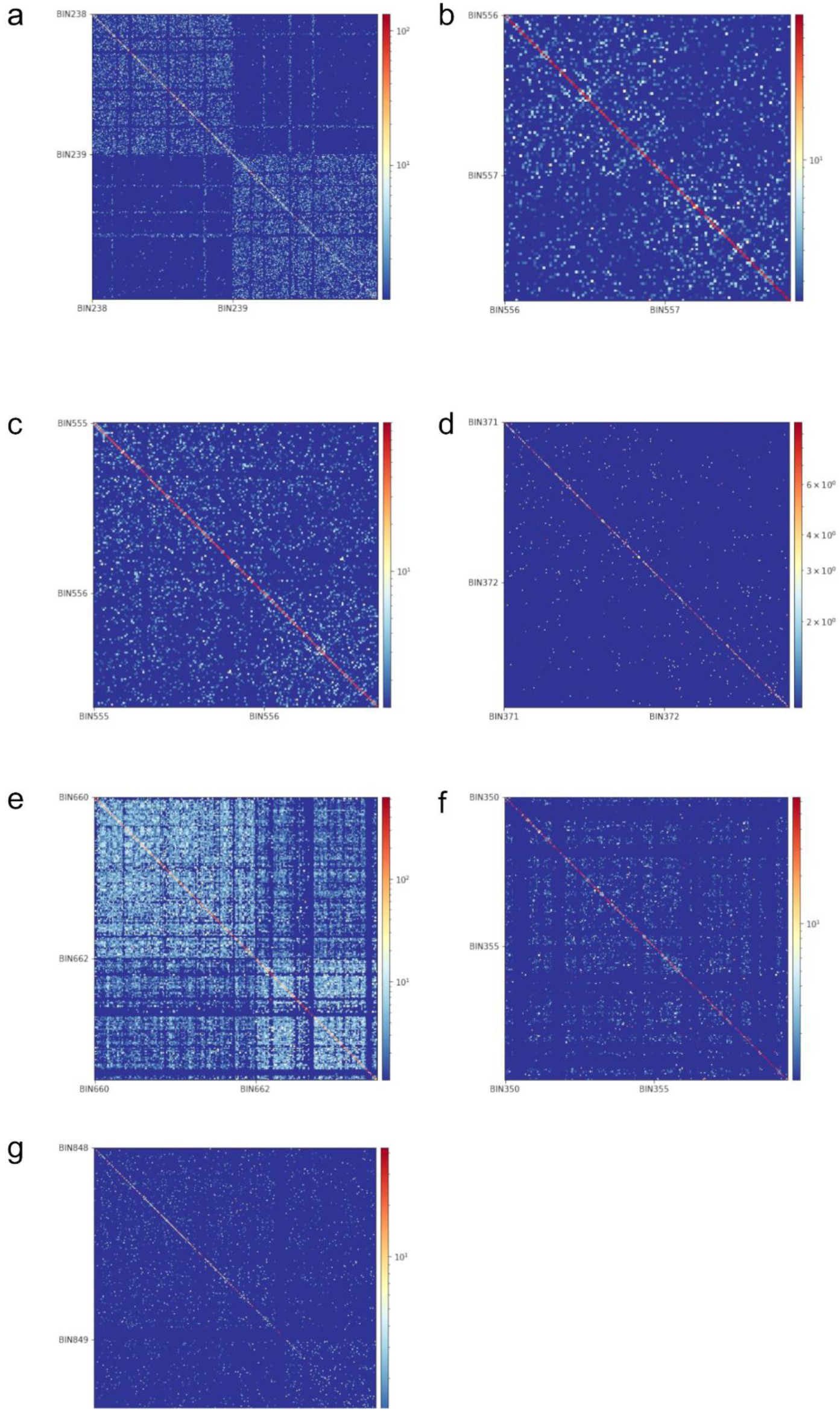
Hi-C contact maps for comparison of HAG pairs. candidate to join at different Nadal Index to support the threshold of Otsu’s method and *k-*means. The different values were (a) 0.15, (b) 0.2, (c) 0.32, (d) 0.35, (e) 0.42, (f) 0.5 and (g) 0.7.

