## Supplementary information for "Eco-evolutionary dynamics of active virus-host interactions in a freshwater lake: revealed through metaHi-C"

**Similarity of linked vHAGs to previously assembled publicly available viral genomes:**
Within linked HAG-vHAGs detected in our metaHi-C dataset, four vHAGs (vHAG0078/0133/42-74/48) were linked to *Ca.* Planktophila representatives with no sequence similarity with the phages inferred for this lineage. However, they showed sequence similarity with six viral sequences within the IMG/VR 4.1 high-confidence database [1] (**Figure SI1**). In addition, HAGs affiliated to genus *Ca.* Nanopelagicus, for which no phages have been reported so far, were linked to four vHAGs (vHAG0841/4531/8307/19061). These vHAGs showed sequence similarity to five phage sequences in the IMG/VR database that lacked any host designation (**Figure SI2**). Active infections were also detected in representatives of the genus *Ca.* Fonsibacter, linked to six vHAGs (vHAG0057/0211/0472/0781/18146/9210). None of these showed similarity to any inferred *Ca.* Fonsibacter phage, like MN692961, but they did share sequence similarity with nine other viral contigs in the IMG/VR database that lack host designation (**Figure SI3**). Similarly, four vHAGs (vHAG0155/6619/11296/26259) were linked to *Ca.* Methylopumilus representatives, with no detectable similarities to phage P19250A, previously isolated as representative of this genus [2]. Additionally, vHAG0155 showed sequence similarity to two phage genomes in the IMG/VR database with no host designation (**Figure SI4**). Three vHAGs (vHAG0048/0141/1242) were linked to *Polynucleobacter* representatives, being the first reported phage genomes infecting this ubiquitous freshwater lineage and shared sequence similarity with 6 phage genomes with no host designation in the IMG/VR database (**Figure SI5**). Additionally, eight vHAGs were linked to *Limnohabitans* representatives and only three (vHAG0007/0123/0467) showed similarities with phages in the IMG/VR database (**Figure SI6**). Our results show that the most abundant and ubiquitous lineages of freshwater bacterial genera are actively infected by viruses and expand the known

diversity of phages infecting these lineages by resolving the host for publicly available phage genomes with no host designation.

**Active metaHi-C derived infections *versus in-silico* inferred virus-host interactions:**

To further evaluate virus-host associations and assess the robustness of the Hi-C-derived linkages, additional *in-silico* host prediction approaches were applied, as detailed below.

(i) Among the 85 vHAGs with Hi-C links to a bacterial HAG, putative host prediction using iPHoP recovered 28 hosts, a 32.9% success rate (**Supplementary Table S5**). Among them, only five predictions (17.8%) matched the Hi-C-based host assignments at the genus level. Nevertheless, discrepancies may reflect broad-host-range infectivity, and the possibility that these viruses infect the taxa designated by iPHoP cannot be ruled out.

(ii) The Actinobacteria-specific transcription factor gene *whiB* [3] was annotated in four of the 18 vHAGs linked to Actinomycetota hosts (vHAG0093 and vHAG0336 to *Streptomyces,* vHAG0823 to *Acidimicrobium* and vHAG42-74 to *Clavibacter*). Nevertheless, the precise virus-host infection pair cannot be extracted from the presence of *whiB* alone.

(iii) CRISPR spacer analysis (cut-off: 100% identity, full-length match) revealed 27 matches involving 26 vHAGs and 16 HAGs (**Supplementary Table S6**). Among these, one virus-host pair was also validated by Hi-C linkage. This vHAG encodes glucosyltransferases, which are thought to mediate DNA glycosylation enabling phages to evade restriction endonucleases and interfere with the CRISPR-Cas system [4] . The detected linkages may also reflect a failed infection event (resulting in a CRISPR spacer acquisition) [5].

(iv) Miniature inverted-repeat transposable elements (MITEs) were also screened to identify virus-host associations [6] . A total of 94 cellular MITEs (cMITEs) were detected in seven HAGs and 16 viral MITEs (vMITEs) in ten vHAGs. However, no matches exceeding 95% identity over the full length were identified. Screening for similar virus-like MITE sequences (si-vMITEs) against the cMITEs database [6] revealed three vHAGs associated with *Pedobacter*, although none were corroborated by Hi-C linkages (**Supplementary Table S7**).

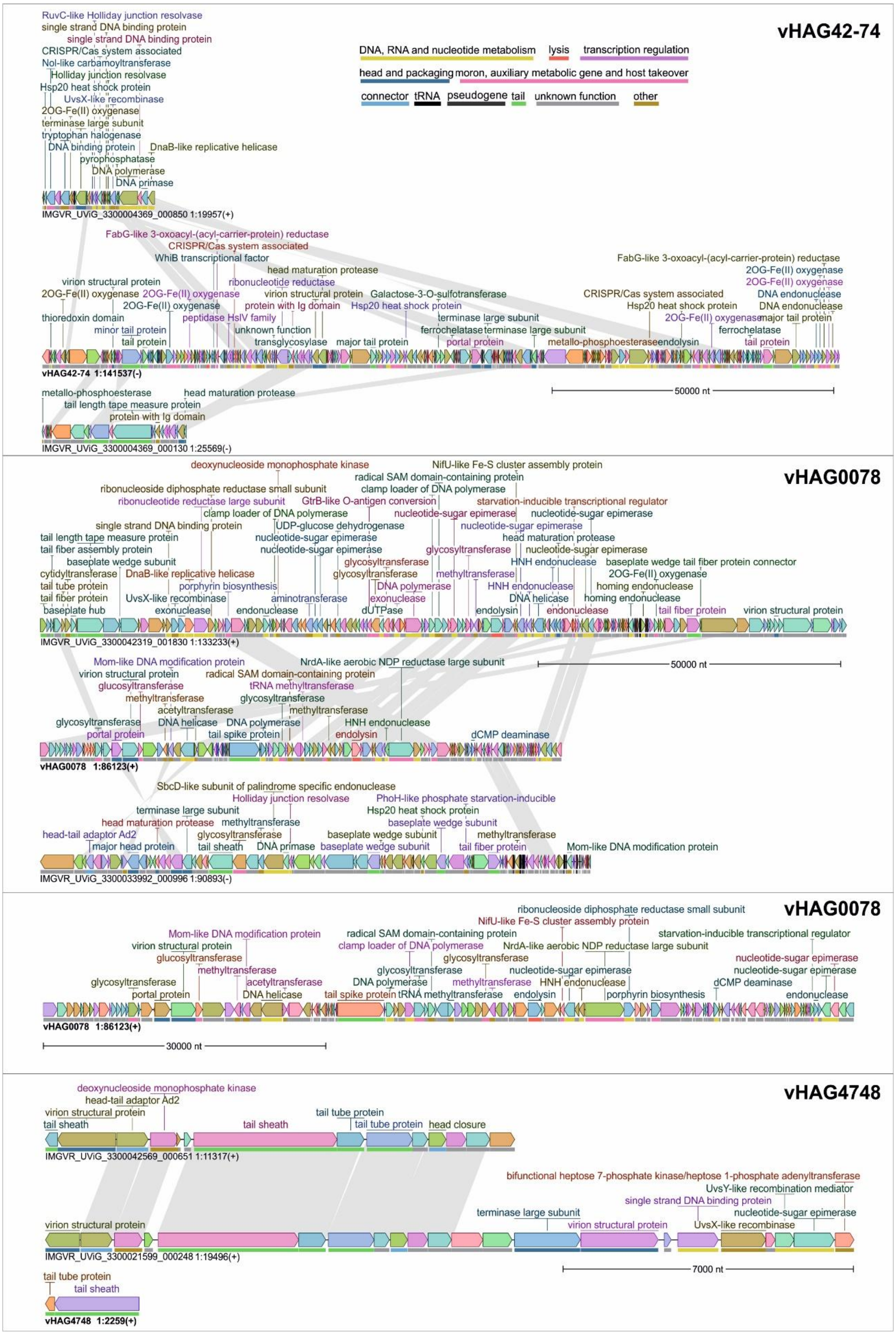


**Figure SI1.** Genomic comparison between *Ca*. Planktophila-associated vHAGs with JGI viral sequences.

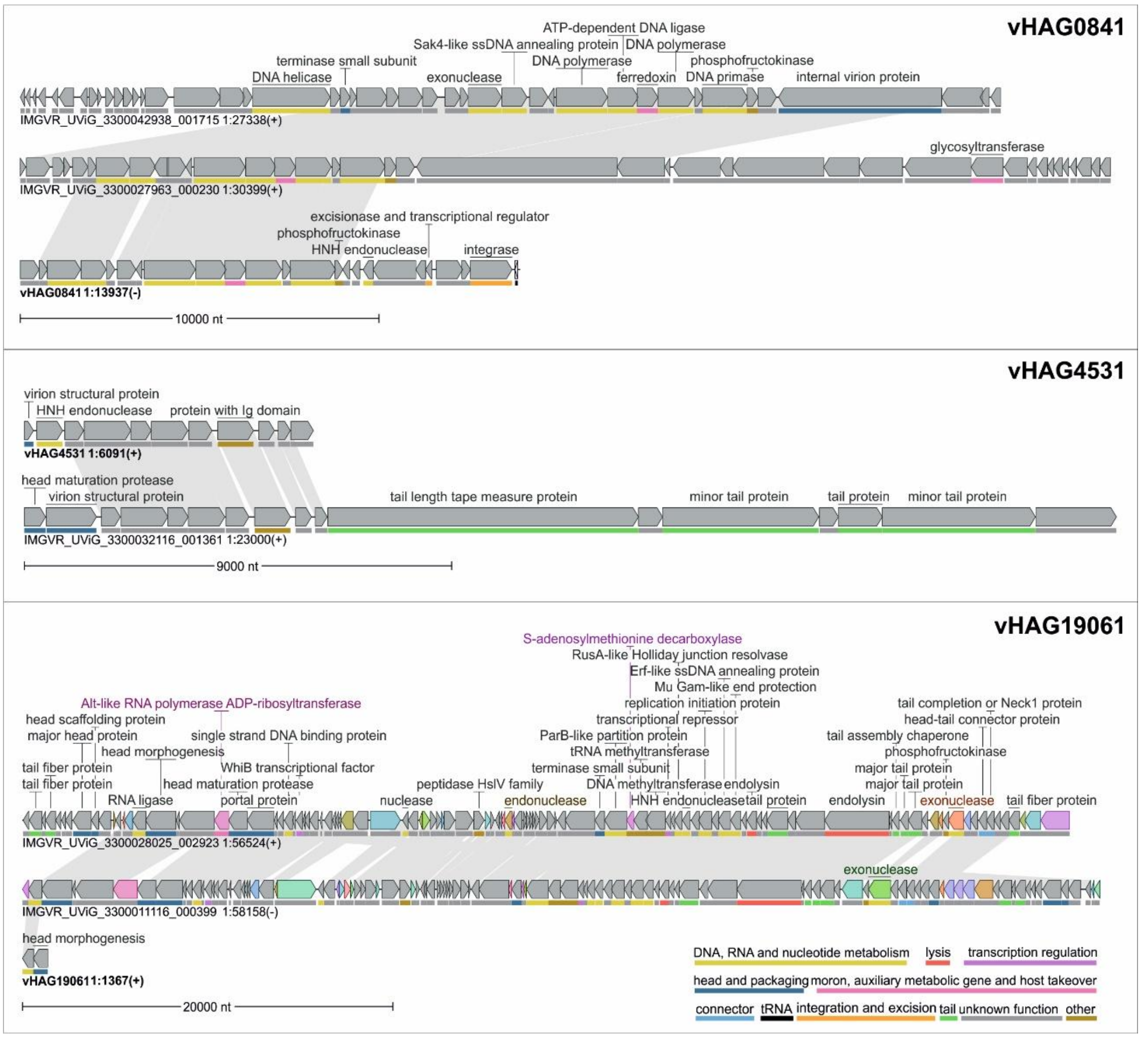


**Figure SI2.** Genomic comparison between *Ca.* Nanopelagicus-associated vHAGs with JGI viral sequences.

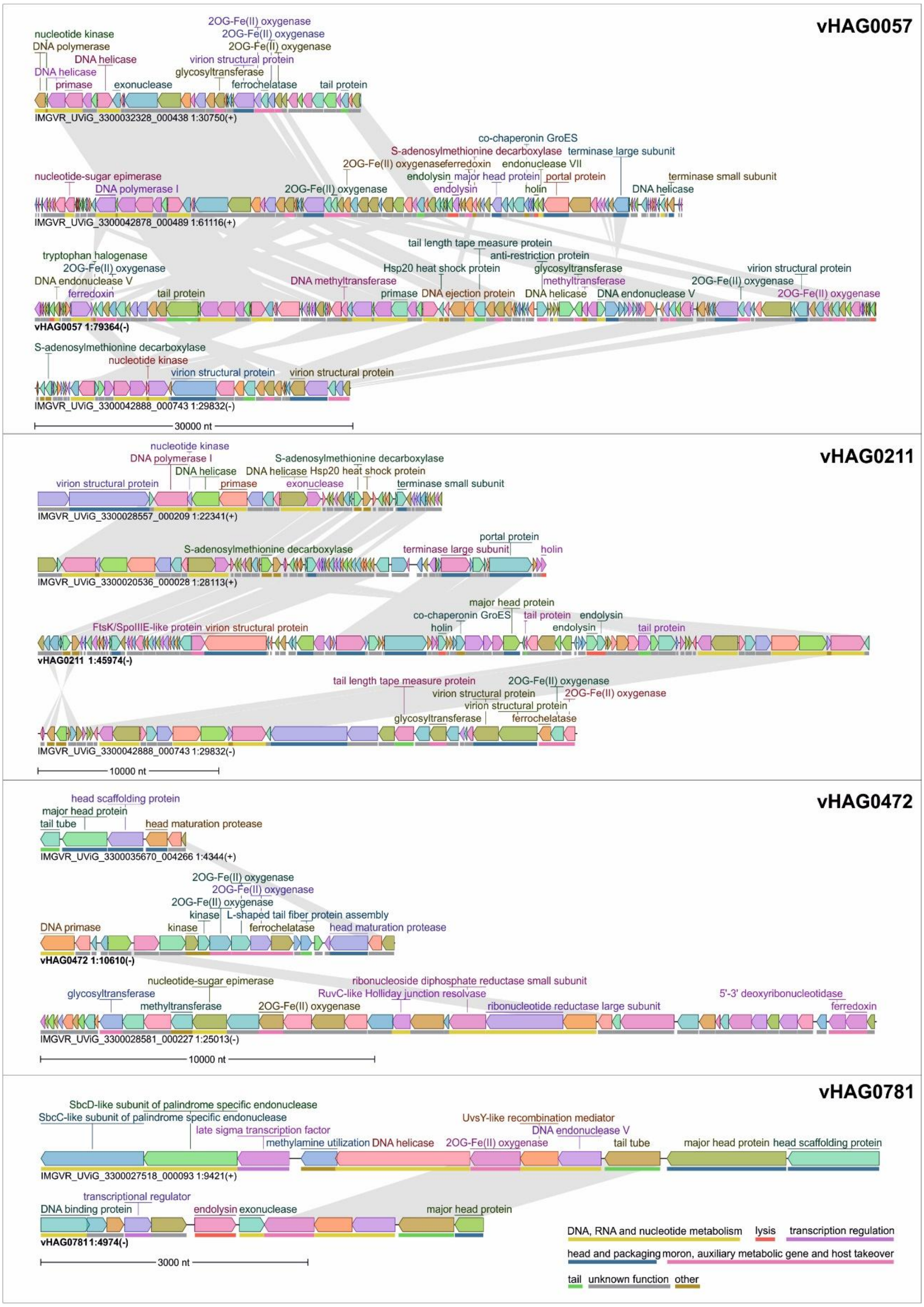


**Figure SI3.** Genomic comparison between *Ca.* Fonsibacter-associated vHAGs with JGI viral sequences.

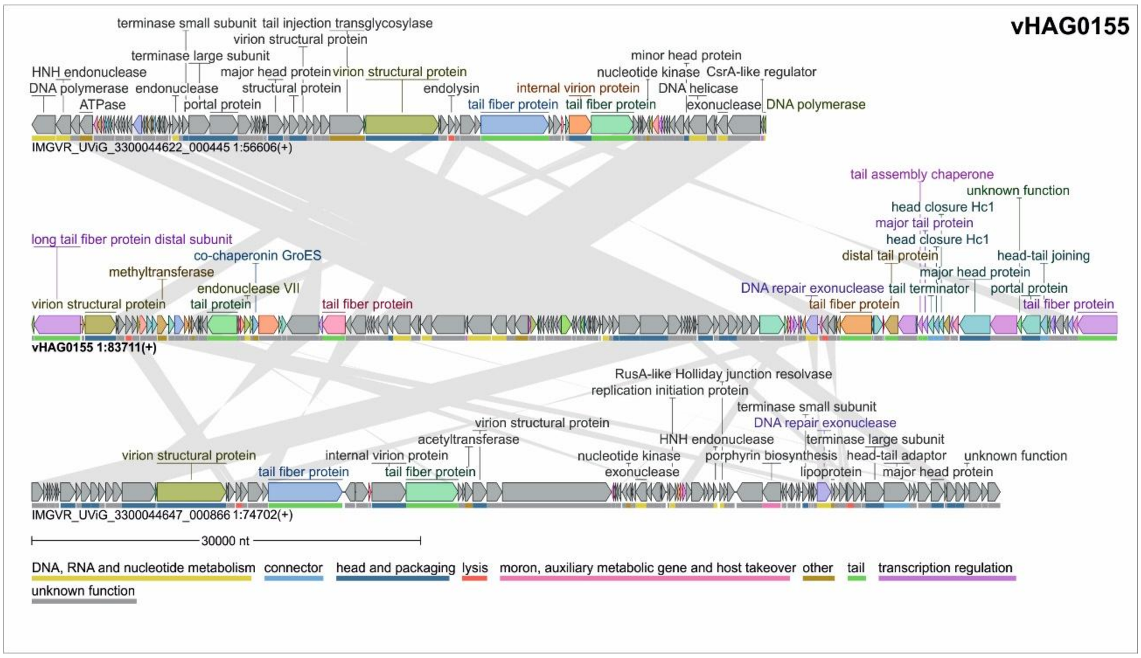


**Figure SI4.** Genomic comparison between *Ca.* Methylopumilus-associated vHAGs with JGI viral sequences.

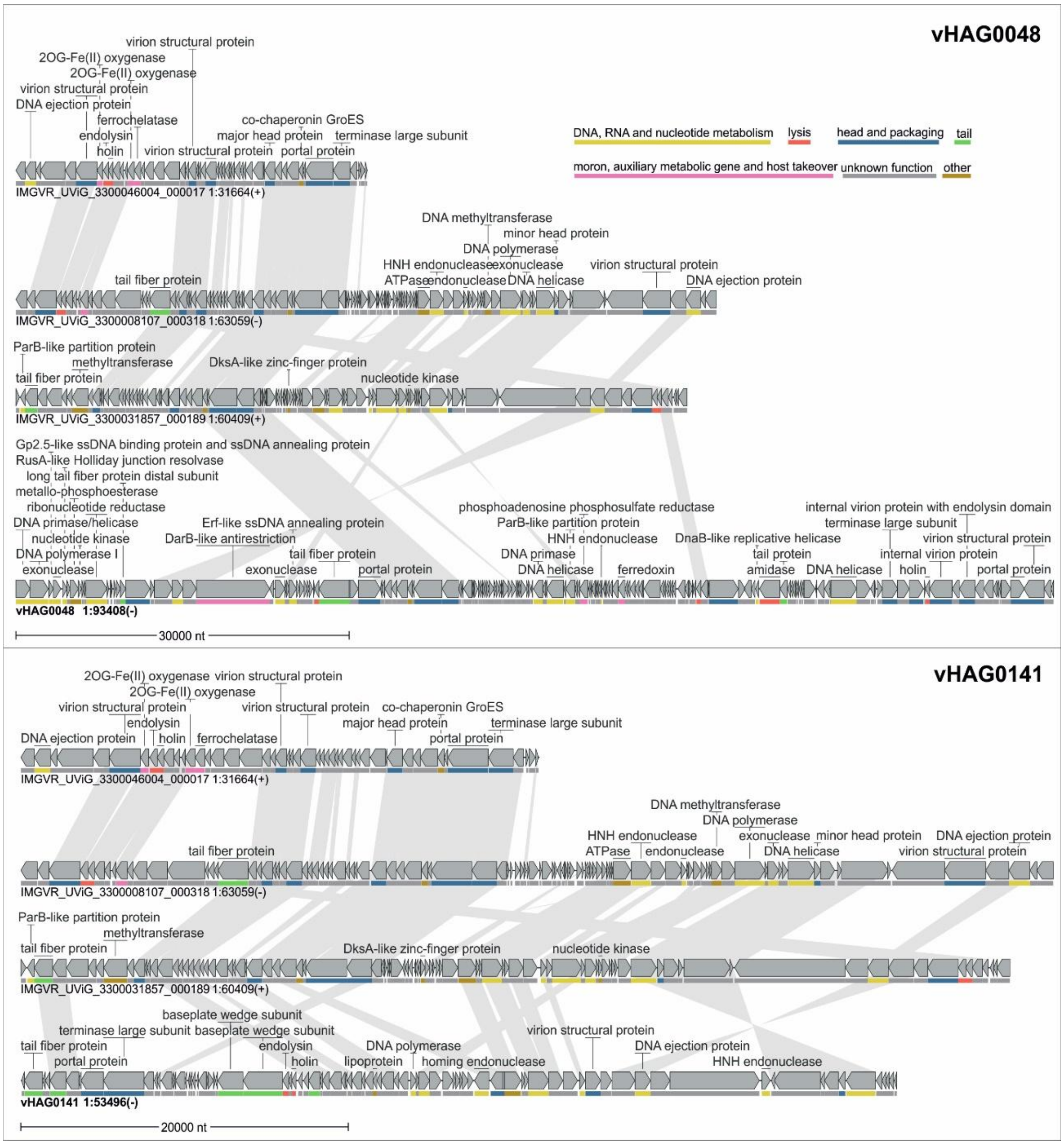


**Figure SI5.** Genomic comparison between *Polynucleobacter*-associated phages with JGI viral sequences.

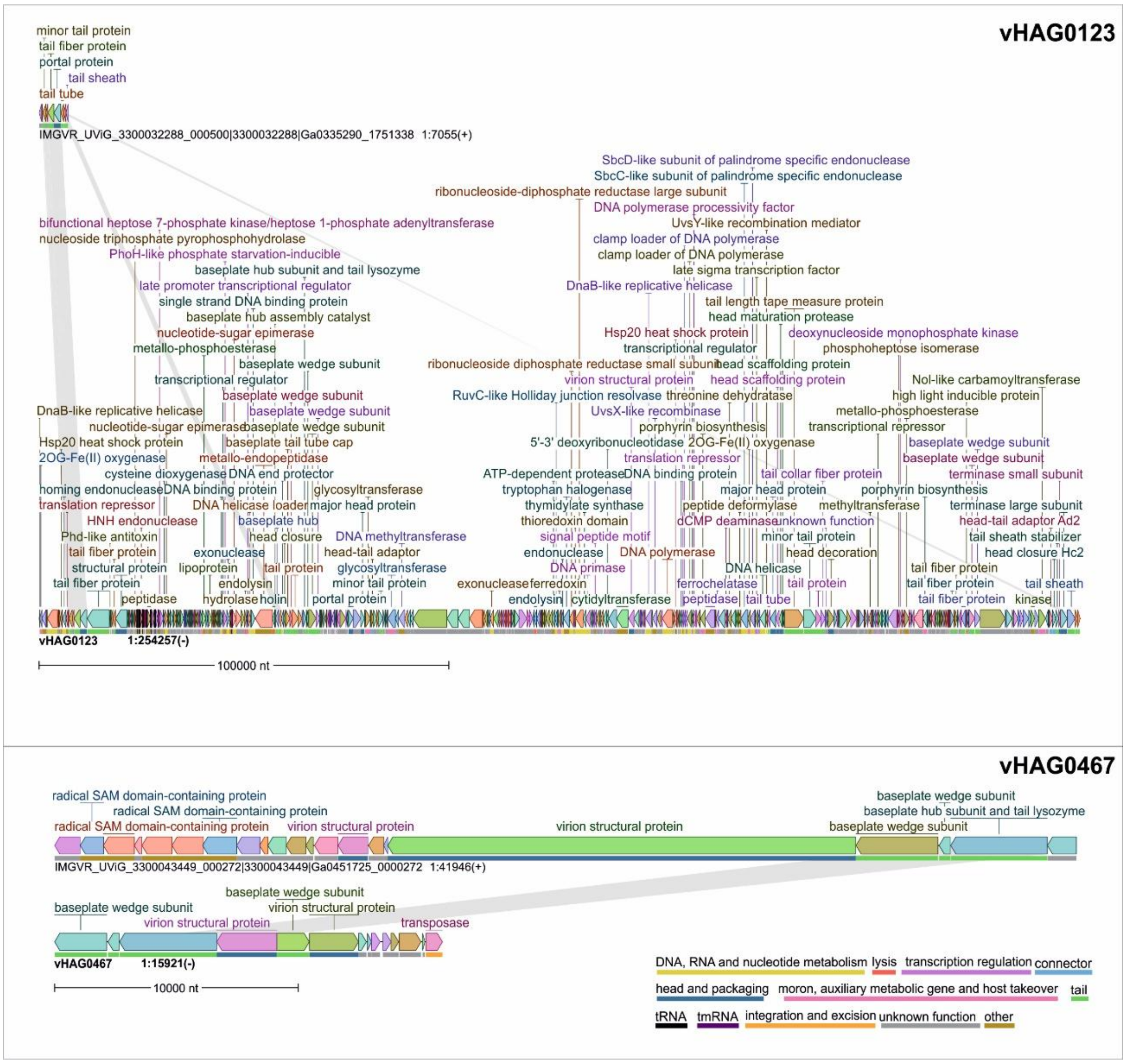


**Figure SI6.** Genomic comparison between *Limnohabitans* associated phages with JGI's viral sequences.
